# Tunable electrostatic interactions of lipid-coated quantum dots with biological membranes

**DOI:** 10.64898/2026.05.21.726631

**Authors:** Lion Morgenstein, Carlos A. Huang-Zhu, Shimon Yudovich, Asaf Grupi, Reid C. Van Lehn, Shimon Weiss

**Affiliations:** Institute for Nanotechnology and Advanced Materials, Bar-Ilan University, Ramat-Gan, 52900, Israel; Department of Physics, Bar-Ilan University, Ramat-Gan, 52900, Israel; Azrieli Faculty of Medicine, Bar-Ilan University, Safed, 1311502, Israel; Department of Chemical and Biological Engineering, University of Wisconsin – Madison, Madison, WI 53706, USA; Department of Molecular and Cell Biology, University of California, Berkeley, Berkeley, California 94720, USA; Department of Chemistry, University of Wisconsin – Madison, Madison, WI 53706, USA; Department of Chemistry and Biochemistry, University of California Los Angeles, Los Angeles, CA 90095, USA; California NanoSystems Institute, University of California Los Angeles, Los Angeles, CA 90095, USA

**Keywords:** Quantum dots, membranes, membrane adsorption, molecular dynamics simulation, FACS

## Abstract

Surface functionalization of inorganic quantum dot nanoparticles is of great interest in the application of these materials toward a wide range of biological applications where membrane interactions are critical. The use of amphiphilic lipids to functionalize the surfaces of quantum dots represents a promising alternative to produce water-soluble and membrane-active materials with facile tuning of the quantum dot’s surface properties. Here, we demonstrate an experimental approach that yields lipid-coated quantum dots with highly tunable surface charge by controlling the concentration of cationic lipids during preparation. Through fluorescence-activated cell sorting assays, we show that these cationic lipid-coated quantum dots can enhance membrane interactions and increase membrane labeling density in live HEK293 cells. We further employed coarse-grained molecular dynamics simulations to model the lipid self-assembly process using an implicit solvent force field and subsequently model the adsorption of lipid-coated quantum dots to model membranes. Our simulations show that we can control the effective surface charge of lipid-coated quantum dots and influence the strength of adsorption to oppositely charged lipid membranes, a process that is mediated by the release of counterions at the quantum dot-membrane interface. This work supports the future development of biocompatible and water-soluble inorganic nanoparticles with highly tunable surfaces, and provides mechanistic insight into how different lipids can influence nanoparticle-membrane interactions at a molecular scale.

## 1. Introduction

Quantum dots (QDs) are semiconductor nanocrystals that are widely employed in biological research as fluorescence imaging tools due to their unique optical properties^1-4^. QDs exhibit several significant advantages over organic dyes. Their tunable core sizes (1–10 nm) allow for a wide range of fluorescence emission peaks with symmetric and narrow spectra. QDs exhibit very broad absorption spectrums, which make them ideal for multiplexed biosensing applications – multiple color QDs can be excited at a single wavelength. Also, QDs can be efficiently excited far from their emission spectra, reducing background and scattering contributions. Moreover, QDs have far greater extinction coefficient, quantum yield, and photostability as compared to their organic dyes counterparts^5^.

Although QDs have many advantages as probes for bioimaging, a drawback is their surface hydrophobicity, which is imparted by nonpolar capping ligands used during synthesis. As a result, QDs need to be functionalized by additional steps in order to render them water-soluble. Multiple functionalization approaches for improved aqueous solubility and colloidal stability have been introduced^6, 7^. One such approach involves ligand exchange, in which the hydrophobic capping surface ligands are replaced by hydrophilic bifunctional ligands^2, 4, 5, 8, 9^. However, ligand exchange should not compromise their optical capabilities: after ligand exchange, QDs need to maintain high photostability as well as enhanced brightness. This is a difficult task since ligand exchange can permanently modify QD surfaces, thereby altering their photophysical properties. Another functionalization approach involves adsorption of amphiphilic molecules such as block copolymers or phospholipid micelles^10-13^ while maintaining the original hydrophobic as-synthesized ligands on the surface. In both approaches (ligand exchange or adsorption), further steps could involve bioconjugation of molecular recognition molecules (such as antibodies, Streptavidin, or other proteins) suitable for fluorescent labeling; such functionalized water-soluble QD constructs are widely available commercially.

Efficient interaction of QDs with biological membranes can offer unique possibilities for cell membrane engineering. One application of membrane-inserted nanocrystals is for optical observation of membrane potential. Voltage-sensitive semiconductor nanoparticles (NPs), displaying the quantum-confined Stark effect (QCSE) ^14^, offer superior properties as membrane potential sensors. Analogous to the atomic Stark effect, the bandgap of semiconductor nanocrystals may be altered as a result of local electric fields inside polarized biological membranes during an action potential, which in turn translates into changes in fluorescence quantum yield, lifetime, and peak emission wavelength^15, 16^. Such QDs could potentially function as a membrane potential sensor^14^. Recently, rod-shaped nanocrystals termed nanorods (NRs) have been used as optical probes for membrane potential^17, 18^. When compared to contemporary organic voltage-sensitive dyes (VSDs), voltage-sensitive QDs have superior characteristics as membrane potential sensors. They (1) have substantially higher voltage sensitivity, (2) are very bright and afford single molecule detection, (3) display a large spectral shift as function of voltage, (4) have a very fast response, and (5) do not bleach, allowing them to be used at very low concentration. Delivery of voltage-sensitive QDs into the cell membrane is a fundamental prerequisite for the successful application of these nanoparticles as membrane potential sensors in biological applications.

Here, we developed a lipid coating technology that controls QD surface properties to promote QD adsorption to the cell membrane. Through the adsorption of lipids onto the surface of the QDs, we engineered particles with high colloidal stability and biocompatibility while tuning their surface charge. The idea of coating the surfaces of inorganic materials with amphiphiles to enable them as membrane-active materials has been previously explored through experiments and molecular dynamics (MD) simulations^19-31^. Such simulations complement experiments by providing insights into molecular-scale interactions that are difficult to visualize through experimental methods (*e*.*g*., localization of charge densities and lipid reorganization within model membranes). In order to control the particle interaction with the membrane, we optimized the lipid-coated QDs (lcQDs) by controlling lipid composition; lipids vary by their hydrophobic alkane chain, which affects their phase transition and fluidity, and their head group, which affects their charge. We chose 1,2-dimyristoyl-*sn-glycero*-3-phosphocholine (DMPC) as the zwitterionic component and 1,2-dioleoyl-3-trimethylammonium-propane (DOTAP) as the cationic component for simplicity and to improve lcQD colloidal stability^7, 32^. We prepared lcQD constructs with varying compositions of lipids per QD particle to control the lcQDs’ effective surface charge through lipid self-assembly. Fluorescence-activated cell sorting (FACS) experiments revealed that the strength of interaction between positively charged lcQDs and HEK293 cells correlates with the initial DOTAP concentration during lcQD preparation. Finally, we used coarse-grained MD simulations to show that positively charged lcQDs adsorb to a negatively charged model membrane composed of 1,2-dioleoyl-*sn-glycero*-3-phosphocholine (DOPC) and 1,2-dioleoyl-*sn-glycero*-3-phosphoglycerol (DOPG) through a local enrichment of anionic lipids and the displacement of salt ions at the NP-membrane interface, providing a molecular mechanism to explain the increased interaction strength observed experimentally.

## 2. Results and Discussion

### Lipid mixture composition controls electrostatic properties of lcQDs

We began our study by tuning the initial lipid composition during lcQD preparation to identify the optimum concentration that achieves full encapsulation of QDs. Briefly, we mixed commercial 5 nm and 11 nm diameter octadecylamine-capped CdSe/ZnS QDs into a 1:1 solution of DMPC:DOTAP at various initial lipid concentrations to control the total number of lipids in solution. Chemical structures and molecular models of the QD and lipids used are shown in Figure 1. An excess of cholate surfactant was added to solubilize the lipids. We measured the ζ-potential of the resulting lcQDs (Figure 2A) to assess their colloidal stability. For both QD sizes, we found that the ζ-potential plateaus above size-dependent lipid-to-particle ratios. We interpreted this plateau as indicating full encapsulation of the QDs and resulting saturation of the QD surface charge since incorporation of additional lipids into the ligand monolayer would reflect a change in the ζ-potential. Based on the conclusions of our previous work,^33^ in which we demonstrated that lipids intercalate into the ligand monolayer to fill free volume, and on the fact that here we add an excess of cholate as a detergent, we assume that multilayer lipid structures^34, 35^ are not formed on the QD surface nor over the self-assembled lipid monolayer. After determining the lipid-to-particle ratio that yields full encapsulation of the QDs (3000 lipids per particle), we further investigated whether the initial mole fraction of DOTAP, *χ*_*DOTAP*_, had an effect on the lcQDs’ effective surface charge. We again prepared lcQD constructs, but this time varied *χ*_*DOTAP*_ from 0.1 to 0.5 and measured their ζ-potentials. Figure 2B shows that increasing the initial concentration of cationic lipids increases the ζ-potential of the lcQDs, indicating a higher positive surface charge and greater stability in solution.

**Figure 1.**
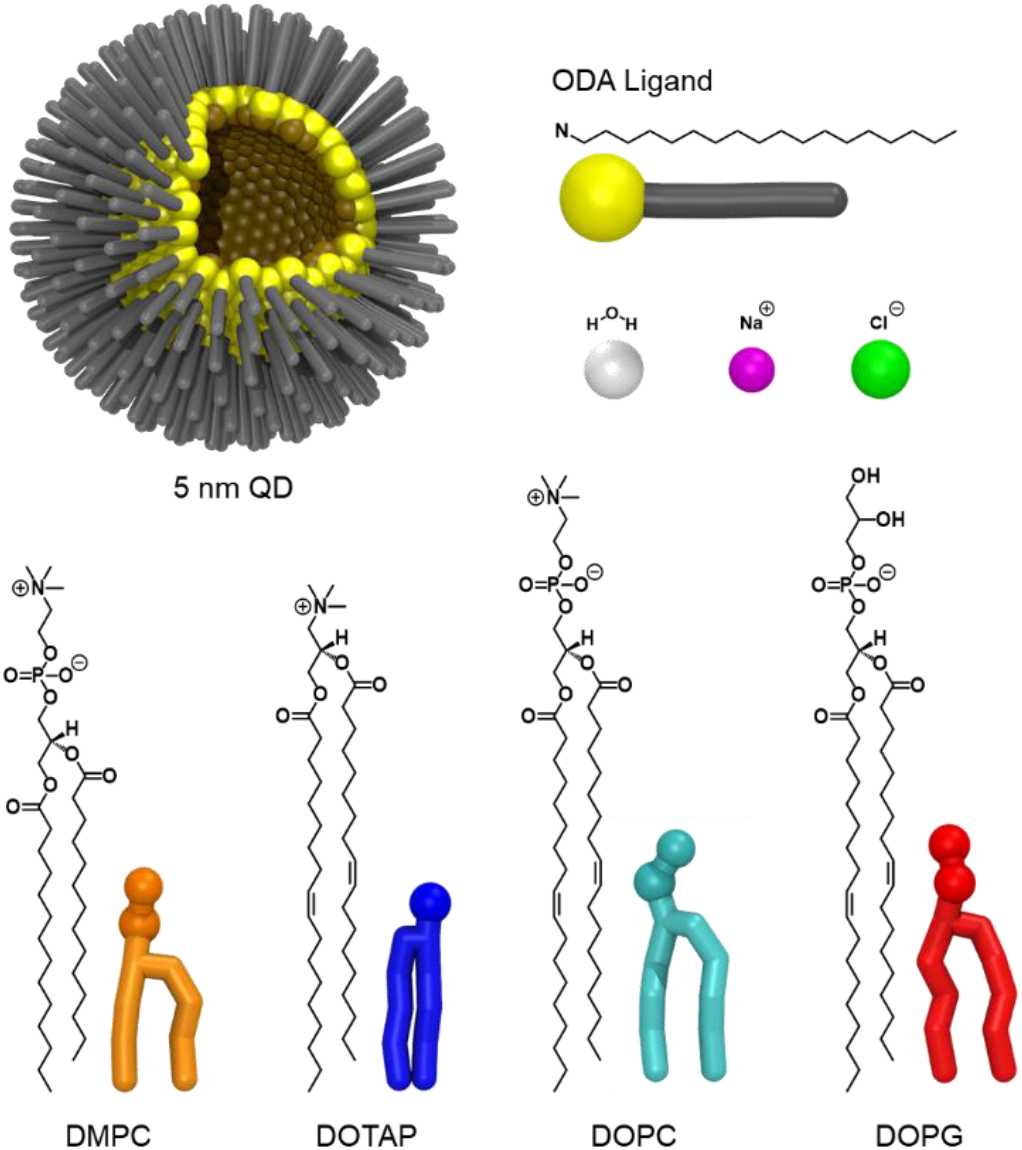
Molecular models used in this study. The CdSe/ZnS QDs capped with octadecylamine (ODA) ligands are colored as follows: CdSe/ZnS shell – tan, ODA propylamine – yellow, ODA alkyl – grey; a portion of the ligands was omitted to show the hollow shell. Lipids are colored as follows: DMPC (modeled as DLPC in the Martini force field) – orange, DOTAP – blue, DOPC – cyan, DOPG – red. Water, sodium, and chloride are colored white, purple, and green, respectively. This color scheme is used throughout the rest of this manuscript.

**Figure 2.**
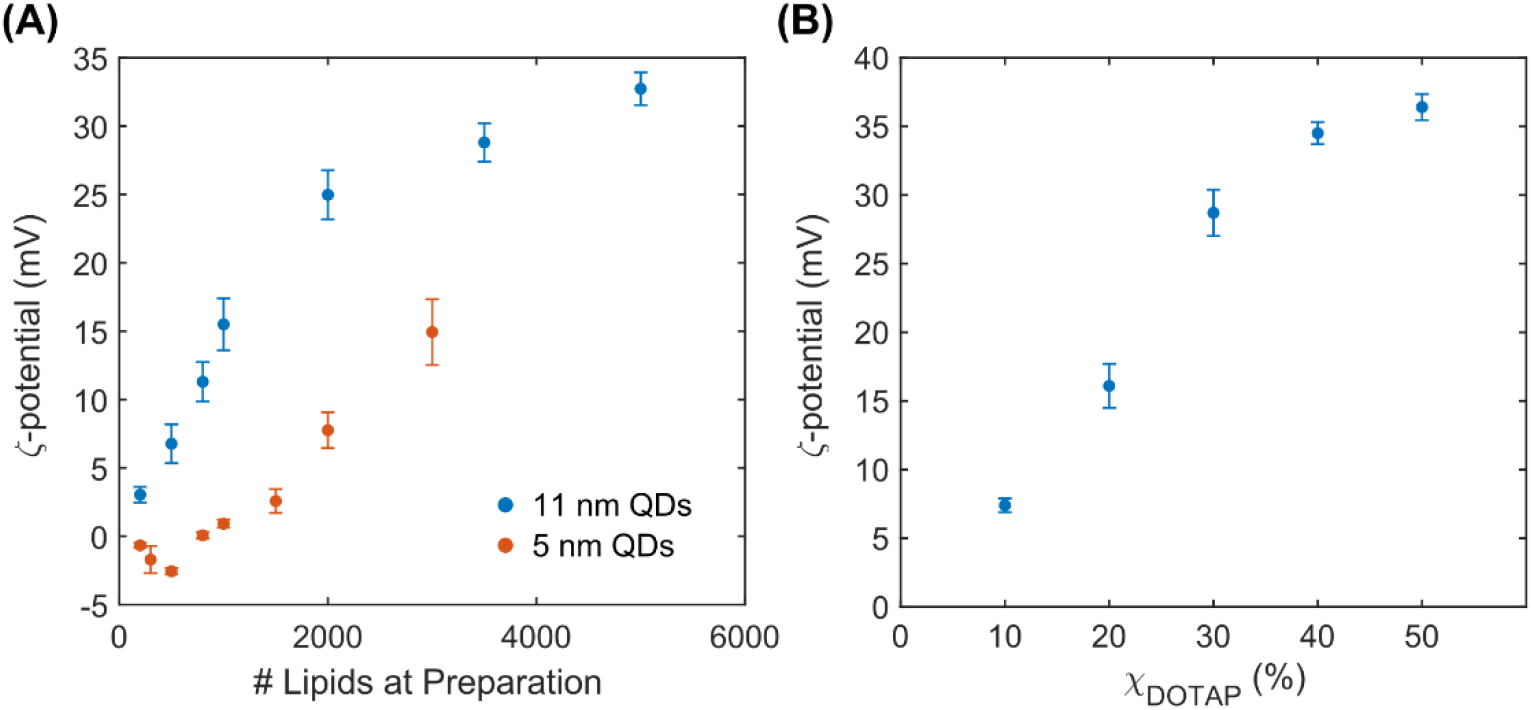
Surface charge dependence of lcQDs on the composition of the encapsulating lipids. (A) ζ-potentials of lcQDs prepared by encapsulating 5 nm (orange) and 11 nm (blue) CdSe/ZnS QDs with varying initial amounts of lipids, spanning the range of 200 to 5000 lipids per particle, using an equimolar mixture of DMPC and DOTAP (*χ*_*DOTAP*_=0.5). (B) ζ-potentials dependence on the initial mole fraction of DOTAP lipids for 11 nm lcQDs, prepared with a total amount of 3000 lipids per QD.

### Self-assembly simulations reveal structures of lipid coatings

To investigate how *χ*_*DOTAP*_ influences the self-assembled lipid coating on the QD, we performed coarse-grained (CG) MD simulations. To achieve the system sizes needed to model lipid self-assembly onto the QD surface, we modeled the self-assembly process using the Dry Martini force field.^36^ Dry Martini is an implicit solvent force field that has interactions parameterized to capture the hydrophobic effect, and has been shown to reproduce related processes, such as vesicle self-assembly^37^ and fusion^38, 39^, in good agreement with more detailed models^40-42^. Since our experiments showed similar trends for the 5 nm and 11 nm diameter QDs, albeit with a larger magnitude for the larger QDs, we chose to model lipid self-assembly on the 5 nm QD to ensure tractable computational expense. The CG model for the ODA-capped CdSe/ZnS QD was taken from our previous work^33^. The computational workflow (schematically shown in Figure S5) mimicked experimental protocols by adding DMPC and DOTAP in a 1:1 ratio into the simulation box to identify the lipid concentration that yields fully encapsulated lcQDs. We then performed simulations with the optimal lipid concentration but varying the initial concentration of DOTAP 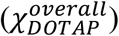. For simplicity and to eliminate NP self-aggregation, we modeled a single QD in the simulation box. The cholate surfactant was not included in the simulations, which is similar to previous molecular simulations that investigated the interactions between lipid-coated NPs and biological membranes under the assumption of a surfactant-free lipid monolayer (*i*.*e*., purely trimethylammonium lipids or phospholipids) and by defining a number of adsorbed lipids *a priori*.^22, 43, 44^ Additional details are provided in the Methods section. Experimentally, the lcQDs are washed after preparation which is expected to readily remove cholate due to its surfactant nature and small molecular size, leaving surfactant-free lipid coatings. To support this assumption, thermogravimetric analysis-mass spectrometry (TGA-MS) was performed to measure the surfactant concentration and showed unmeasurable trace amounts (data not shown).

Figure 3A shows that the fraction of DOTAP in the simulated surface lipid coating after self-assembly 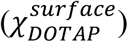 does not change as the total number of lipids in the mixture increases. This invariance indicates that the initial concentration of cationic lipids indeed determines the effective surface charge of the lcQDs. We show that adding 500 total lipids yields partially encapsulated lcQDs, while fully encapsulated particles are observed at 1,500 total lipids and above. Encapsulation was assessed by quantifying the total number of lipids that composed the lipid coating in Figure S6 along with the solvent-accessible surface area (SASA) of the QD in Figure S7. Even though the QD’s average SASA decreased as the total number of lipids increased, indicating encapsulation, at significantly high lipid concentrations inconsistent lipid coatings were observed (simulation snapshots in Figure 3A). When the self-assembly process was initiated with at least 2,000 total lipids, we obtained lcQDs coated with multilayer lipid structures. Visual inspection (Figure S8) revealed that the multilayer structures are similar to vesicle hemifusion states^34, 35^ where the lipid coating partly fuses with a vesicle only through the outer leaflet and forms a hydrophilic pocket. These structures are expected to be high energy states because of the hydrophilic pocket and the lipid line tension formed by the irregular lipid arrangement. Further inspection revealed that the first step leading to the formation of these multilayer structures is the fusion of a vesicle that self-assembles in solution with the lcQD; the size of the then dictates the shape of the lcQD construct (*e*.*g*., large vesicles lead to the lcQD from the 3,500 total lipids simulation shown in Figure 3A). These structures are not surprising due to the high lipid concentration. However, during the experiments, the formation of multilayer structures was addressed and eliminated by the addition of cholate to solubilize vesicles and a filtration step to remove any multilayer structure that may have been formed. Based on these observations, from our simulations we chose lcQDs that did not have these multilayer structures for subsequent analyses.

**Figure 3.**
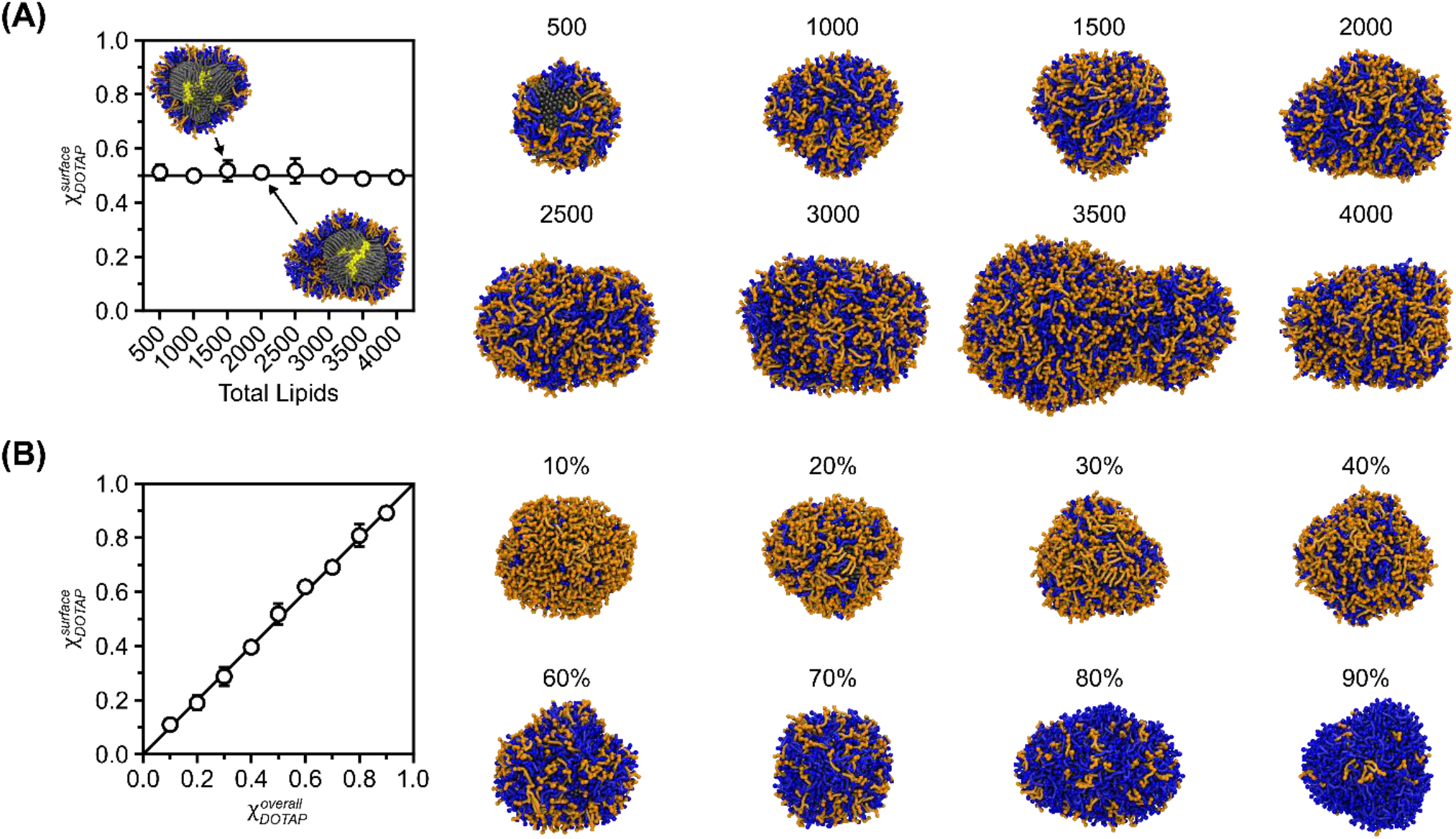
Simulated encapsulation efficiency and surface charge of lcQDs is controlled by initial lipid compositions. (A) Fraction of DOTAP in the lipid coating as a function of the total number of lipids in solution and representative snapshots of the resulting lipid-coated QDs. (B) Fraction of DOTAP in the lipid coating as a function of the DOTAP concentration in solution and representative simulation snapshots of the resulting lipid-coated QDs. All simulation snapshots in the manuscript were rendered using the Visual Molecular Dynamics software^45^. The error in (A) and (B) was computed as the standard deviation across four replicates.

After identifying the total number of lipids per particle that consistently yields fully encapsulated lcQDs and fewer multilayer structures (1,500 lipids per particle) from Figure 3A, we screened the effects of varying 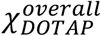 in Figure 3B. We found that 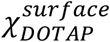 varies linearly with 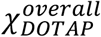, in agreement with our previous hypothesis from experimental observations and Figure 3A. We thus refer to these quantities simply as *χ*_*DOTAP*_ in the following sections. Simulation snapshots reveal that lipids intercalate into the ligand monolayer to ensure hydrophobic voids are filled with lipid tails (Figure 3B and Figure S9). After these voids are filled, additional lipids adsorb over the remaining exposed alkyl chains to ensure full encapsulation, and in the process, lipid tails become intercalated within the lipid coating in an interlocked fashion to prevent their desorption (Figure S8-S9). While the experimental ζ-potential appears to deviate from linearity at a high value of *χ*_*DOTAP*_, the encouraging agreement between experimental and computational trends suggest that the simulated structural organization of lipids on the lcQD surface is reasonable and that overall surface charge of lcQDs is controlled by tuning the initial concentration of lipids during the preparation step. In the following sections, we thus present data to study mechanisms of adsorption for positively charged lcQDs, under the assumption that the fully saturated lcQDs from experiments are equivalent to the simulated lcQDs with regard to lipid coating composition and lipid arrangement.

### Lipid coating dictates interactions between lcQDs and HEK293 cells

To assess the effect of *χ*_*DOTAP*_ on the interaction between lcQDs and biological membranes, we performed FACS experiments using HEK293 cells. Since these cells have an anionic outer membrane leaflet^46-48^, we hypothesized that the lcQDs’ overall surface charge would affect the interaction strength with the cell membrane, with larger lcQD surface charge leading to increased membrane adsorption. Cells were incubated with a series of 11nm lcQDs of varying surface charge but constant lipids per particle ratio, above the encapsulation ratio, all diluted to the same concentration, and washed with phosphate-buffered saline (PBS) buffer. In these experiments, lcQDs that interact strongly with the cell membrane were expected to remain adsorbed, leading to increased fluorescent signal per cell. The lcQD-labeled cells were run through a FACS instrument, and the single-cell fluorescence intensity at the QDs’ emission spectral range was measured. Figures 4A and 4B show FACS histograms and mean fluorescence emission intensity, respectively, from lcQD-labeled cells, performed on 10,000 cells for each lcQD construct. The measured intensities in Figure 4A reveal an increase of up to four orders of magnitude compared to unstained cells as a function of the lcQDs’ ζ-potential. Because cell staining was performed in PBS, which contains ∼140 mM NaCl, these results indicate that the interaction of lcQDs with the membrane increases with ζ-potential, despite electrostatic screening at biophysically relevant salt concentrations. Thus, by simply tuning the initial lipid composition, one can control the macroscopic properties of lcQDs and consequently their interactions with biological membranes. Finally, we note that increasing the lcQDs’ surface charge (*i*.*e*., by increasing *χ*_*DOTAP*_) resulted in decreased quantum yield, which corresponds to a previously-described photophysical effect^49, 50^. To assess the relative brightness, we divided the fluorescence signal by the optical density (O.D.) and normalized the results against the brightest construct (Figure S2). Figure S3 shows the recalculated results of Figure 4A with this correction. Considering that the reduction in emission occurs for lcQDs with large ζ-potentials, one can suggest that the actual degree of membrane labeling for such particles is even higher than indicated by Fig. 4A. This correction allows for even better separation of the corresponding FACS histograms, resulting in improved differentiation between lcQD samples of different surface charge.

**Figure 4.**
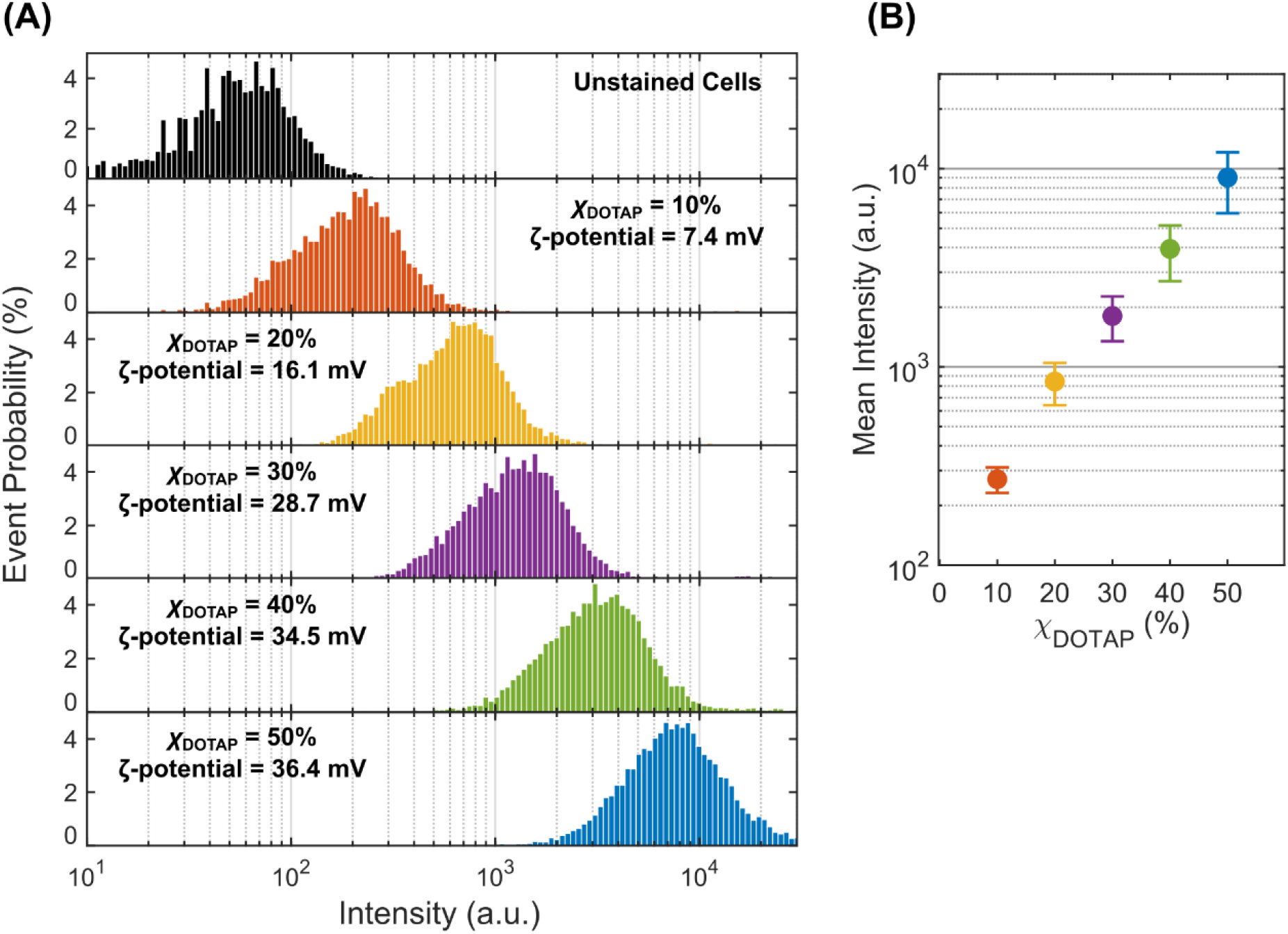
Dependence of live-cell lcQD labeling density on encapsulating lipids charge. (A) and (B) show fluorescence-activated cell sorting (FACS) cell population histograms and mean emission intensity, respectively, of HEK293 cells stained with a series of lcQDs of varying surface charge. All lcQD constructs were encapsulated with 3000 lipids per particle, composed of a DMPC:DOTAP lipid mixture containing a molar fraction of DOTAP varying from 10% to 50%.

### Simulations reveal unbiased lcQD adsorption to anionic membranes

We sought to provide mechanistic insight into how the surface lipid composition of lcQDs can amplify the staining intensity and efficiency, particularly at the salt concentrations studied. To better capture electrostatic interactions compared to the self-assembly simulations, we performed simulations of the lcQDs and a model lipid membrane (80% zwitterionic DOPC and 20% anionic DOPG) using an explicit solvent coarse-grained force field, Martini 2.3^51^, using the refined polarizable water model^52^. These simulations were performed at physiological salt concentrations to screen the electrostatic interactions between charged lipids which are essential in modulating initial lcQD adsorption and capturing relevant biomolecular interactions. Since using the 5 nm diameter QDs from Figure 3B to perform these simulations would require prohibitively large systems sizes, we decided to use smaller 2 nm diameter QDs based on our previous observation that larger QDs exhibit qualitatively consistent trends in lipid encapsulation efficiency. Self-assembly simulations yielded equivalent results for these QDs (Figure S10-S12) from which we extracted lcQDs at three different values of *χ*_*DOTAP*_ (0.1, 0.5, and 0.9) that are representative of the range of DOTAP compositions.

Based on a preliminary set of simulations that showed variations in unbiased adsorption (*e*.*g*., lcQDs would either partition to the membrane surface or into solution), we designed our simulation protocol around the quantification of unbiased adsorption events. Briefly, we placed each lcQD randomly oriented at a minimum distance of approximately 2 nm from the surface of an 18 nm × 18 nm membrane. The system was solvated with Martini polarizable water^53^ and 150 mM NaCl; counterions were also added to neutralize the effective charge of the lcQD. For each of the lcQDs with *χ*_*DOTAP*_ of 0.1, 0.5, and 0.9 we performed 16 × 50 ns unbiased simulations. This large number of unbiased simulations permitted the lcQDs to sample a variety of local membrane environments due to differences in initial configurations. Additional details are provided in the Methods section.

For each replicate of each set of simulations, we computed the minimum distance between any lcQD bead and any membrane lipid bead throughout the simulation and observed large variations that reflected how strongly the lcQD interacts with the lipid membrane (Figures S14-S16). Figure 5 shows representative behaviors in the limits of weak and strong adsorption. Large fluctuations in the minimum distance as a function of simulation time indicate that the lcQD moves back and forth between the bulk solvent and the membrane interface, whereas small fluctuations within a minimum distance of 0.5 nm to 1.0 nm corresponds to the lcQD fluctuating at the membrane interface. In the latter case, there are weak interactions between lipids from the lipid coating and those from the membrane, as shown in Figure 5A, which slightly deform the membrane, resulting in an average minimum distance of 0.50 nm over the last 5 ns for the simulation corresponding to Figure 5A. The deformation of the membrane is apparent from the plot of the lipid number density, which shows pronounced negative curvature in the vicinity of the lcQD although with no notable lipid enrichment (*i*.*e*., no increased local lipid density). This state reflects only a few positively charged DOTAP headgroups on the lcQD surface coordinating with the negatively charged headgroups from the membrane’s DOPG lipids. This behavior contrasts with Figure 5B which shows DOTAP headgroups fully embedded within the headgroup region of the membrane, resulting in an average minimum distance of 0.44 nm over the last 5 ns of the corresponding simulation. This intercalation is mediated by anionic DOPG lipids being localized at the lcQD-membrane interface and coordinating with the cationic DOTAP lipids, as shown by the local enrichment of lipid beads in the number density plot and without membrane deformation being induced.

**Figure 5.**
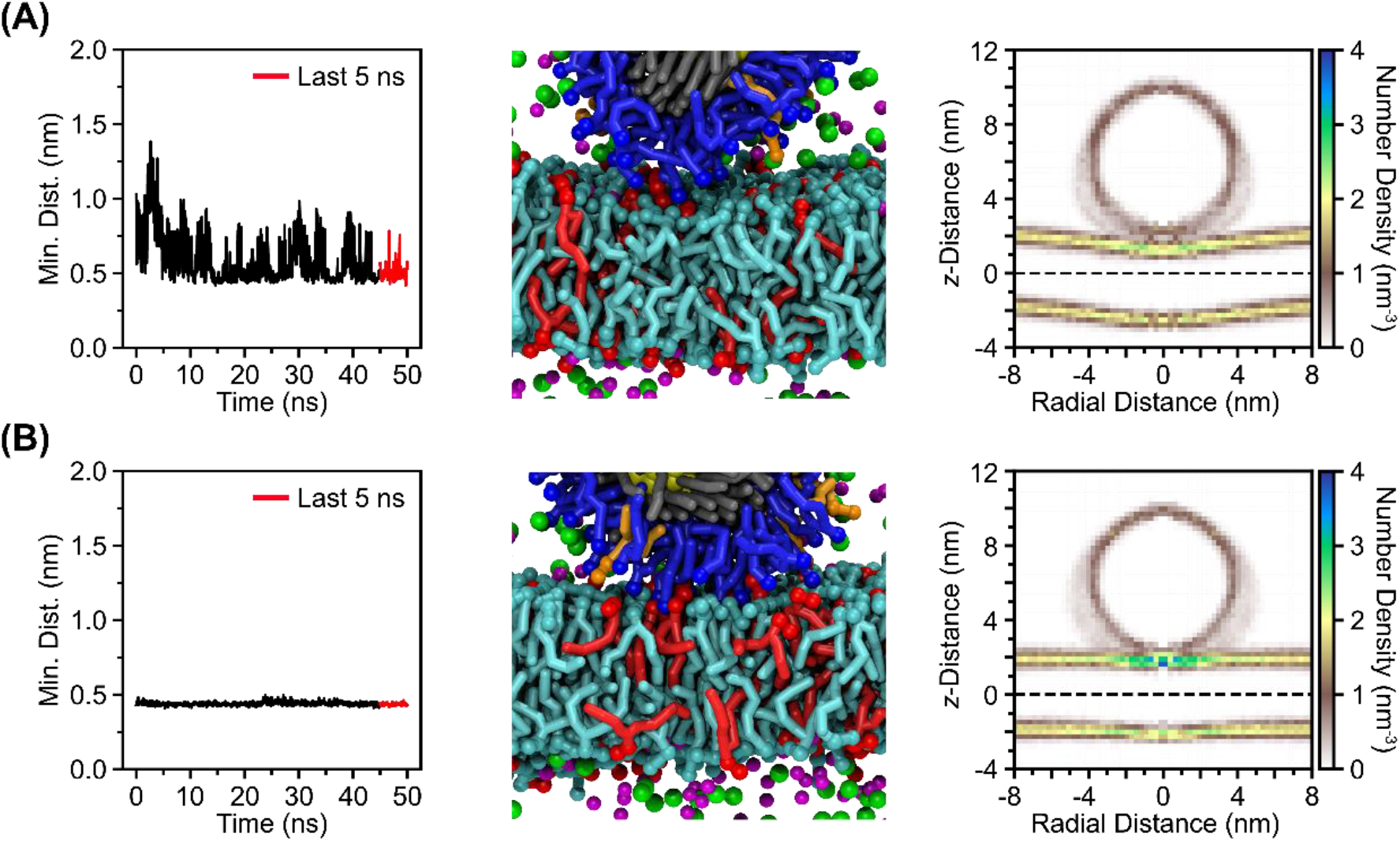
Minimum distance between lcQD and membrane correlates to local lipid enrichment. (A) Replicate in which weak adsorption is observed, and negative membrane curvature is induced. (B) Replicate in which strong adsorption is observed with a relaxed membrane and a local enrichment of lipid headgroups at the lcQD-membrane interface. Data correspond to replicates with *χ*_*DOTAP*_ of 0.9. Number density plots were generated using the phosphate headgroup of DMPC, DOPC, and DOPG, and the choline headgroup of DOTAP. Bins were 0.26 nm in the radial dimension and 0.26 nm in the z-dimension, with the coordinate system centered on the lcQD. Water CG beads were omitted from the simulation snapshots for clarity.

### Increasing DOTAP concentration promotes stronger membrane adsorption

Based on the variations of the minimum distance, we established cutoff values to determine whether a particular simulation replicate resulted in no membrane adsorption, weak adsorption, or strong adsorption (Figure 6A). Based on the number density plots, and the average minimum distance over the last 5 ns of simulation for each replicate, we identified that an average minimum distance between 0.50 and 0.90 nm was consistent with weak adsorption of the lcQD to the membrane. An average minimum distance greater than 0.90 nm was classified as no adsorption and a minimum distance less than 0.50 nm was classified as strong adsorption. Each replicate was classified according to these cutoff values, and their probability was computed in Figure 6B.

**Figure 6.**
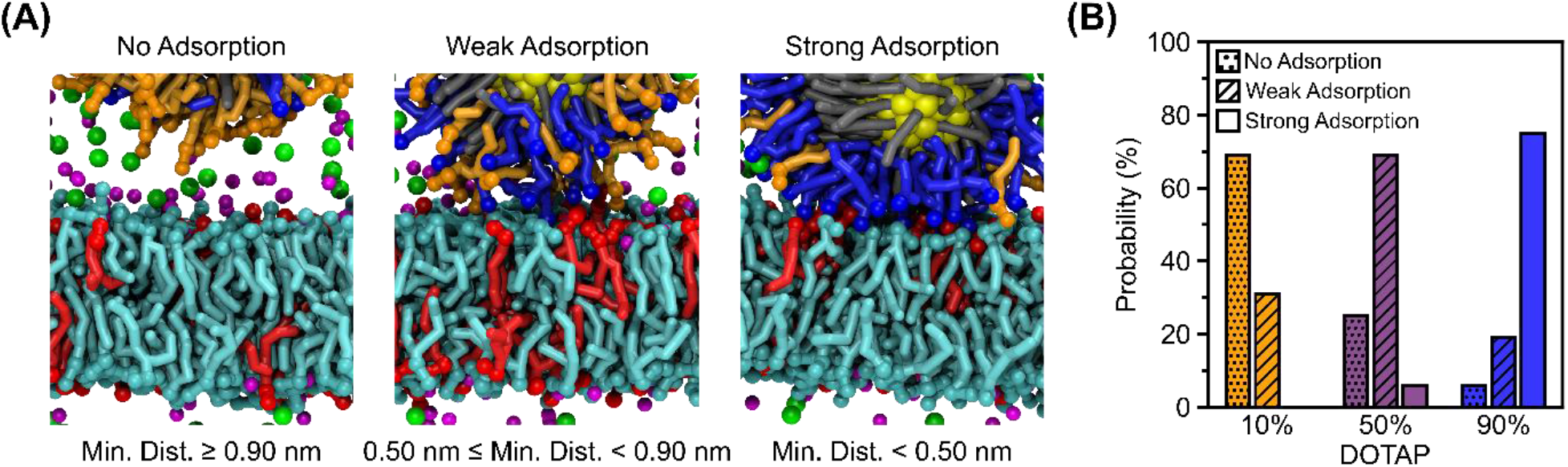
Effective surface charge of lcQDs determines strength of adsorption. (A) Simulation snapshots representative of the three adsorbed states. (B) Probability of observing no adsorption, weak adsorption, or strong adsorption within the ensemble of simulations for lcQDs with *χ*_*DOTAP*_ of 0.1, 0.5, and 0.9 (16 simulations in each ensemble). Water CG beads were omitted from the simulation snapshots for clarity.

For lcQDs with *χ*_*DOTAP*_ = 0.1, 69% of the replicates resulted in no adsorption and the remaining 31% resulted in weak adsorption; no replicate showed strong adsorption. For lcQDs with *χ*_*DOTAP*_ = 0.5, only 25% of the replicates resulted in no adsorption; 69% of the replicates showed weak adsorption and the remaining 6% resulted in strong adsorption. Finally, for lcQDs with *χ*_*DOTAP*_ = 0.9, only 6% of the replicates showed no adsorption and 19% showed weak adsorption; the remaining 75% showed strong adsorption. These trends generally agree with our experimental FACS measurements (Figure 4A) by showing that increasing the effective surface charge of the cationic lcQDs promotes stronger adsorption to the anionic membrane. The calculated probabilities for each state (Figure 6B) relative to each other are also in agreement of our hypotheses. The lcQDs with *χ*_*DOTAP*_ = 0.1 showed no replicate where the QDs adsorbed strongly, and the opposite is observed for lcQDs with *χ*_*DOTAP*_ = 0.9 where only once did the lcQD partition into solution.

### Lipid-lipid interactions and displacement of ions drive lcQD adsorption

Since Figure 6 suggests electrostatic-mediated adsorption of lcQDs despite the relatively high salt concentration (150 mM) which is expected to screen these interactions, we further investigated potential descriptors that may be driving this process. In Figure 7 we quantified the number of charged species within the NP-membrane interface for each of the most probable adsorbed states of Figure 6B (*i*.*e*., “no adsorption” for the ensemble of simulations with *χ*_*DOTAP*_ = 0.1, “weak adsorption” for *χ*_*DOTAP*_ = 0.5, and “strong adsorption” for *χ*_*DOTAP*_ = 0.9). This approach ensures that each state is described with accurate statistics by averaging over multiple independent trajectories. The interface region was defined as the cylindrical region between the center of mass of the QD and the midplane of the membrane with a radius equal to the average effective radius of the lipid coating headgroups relative to the center of the QD, as shown schematically in Figure S13.

**Figure 7.**
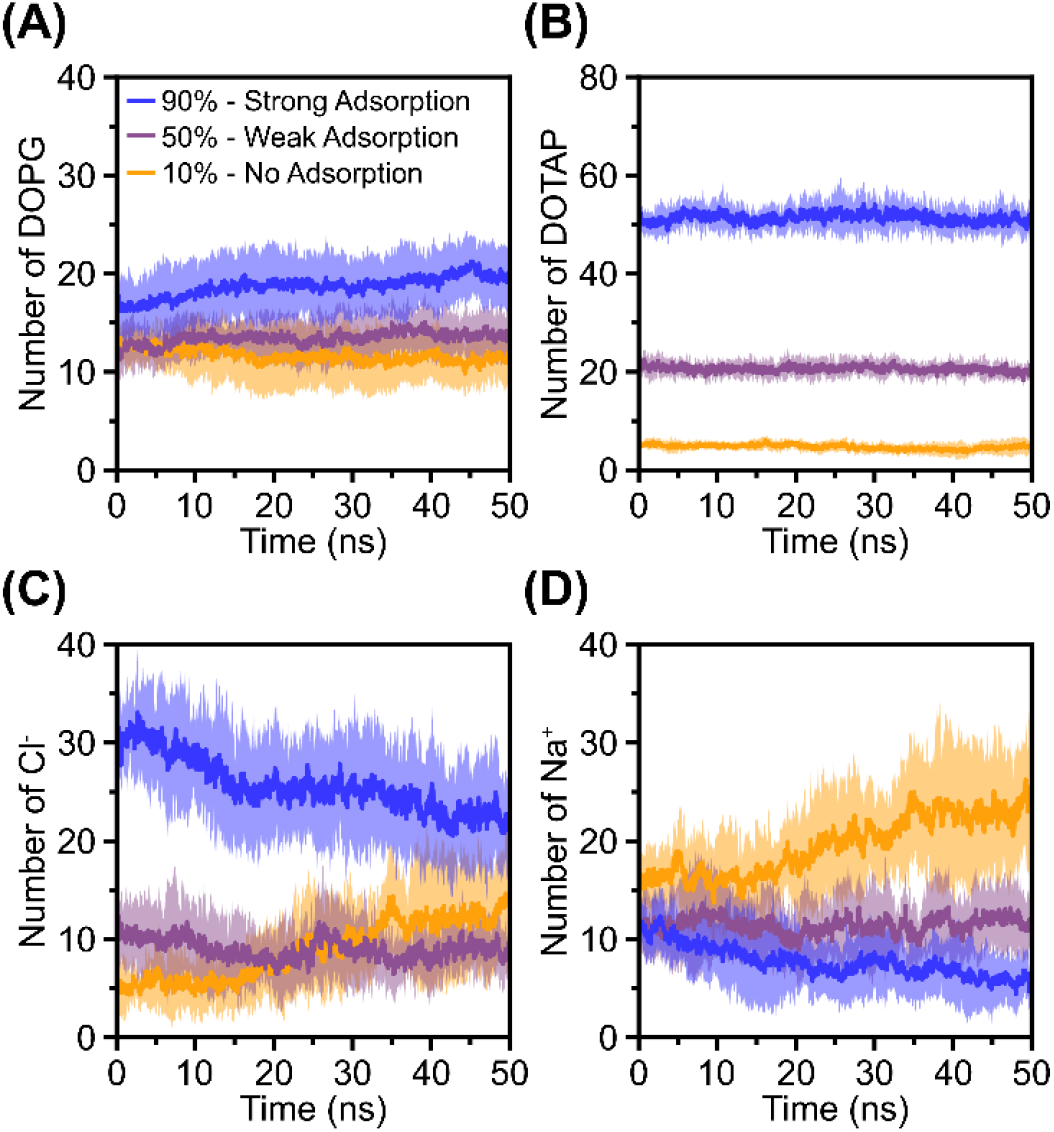
Average number of charged species within the cylindrical region centered at the NP-membrane interface. The error was computed as the standard deviation within each ensemble of simulations (*i*.*e*., 11 replicates for lcQDs with *χ*_*DOTAP*_ of 10% or 50% and 12 replicates for lcQDs with *χ*_*DOTAP*_ of 90%).

Figure 7A shows that DOPG lipids in the membrane reorganize over the simulation timescale in order to increase the DOPG density at the lcQD-membrane interface to enable strong adsorption for the lcQDs with *χ*_*DOTAP*_ = 0.9; this enrichment is not observed for the other two lcQDs. This observation indicates that favorable DOPG-DOTAP electrostatic interactions drive changes to lipid organization at the lcQD-membrane interface. However, DOTAP rearrangement within the lipid coating is negligible for all lcQDs as suggested by Figure 7B, which indicates that lipid mobility is restricted due to the intercalation of lipid tails within the ligand monolayer on the QD. For the lcQDs with *χ*_*DOTAP*_ = 0.9, the enrichment of anionic lipids at the lcQD-membrane interface leads to a corresponding displacement of chloride and sodium ions as shown in Figures 7C and 7D. The data in Figure 7C show first that the number of chloride ions follows trends in the DOTAP content, with higher densities initially present in the region between the lcQD and membrane for higher DOTAP content since the chloride ions compensate for the cationic DOTAP charge. The number of chloride ions substantially decreases for the strongly adsorbing lcQDs with *χ*_*DOTAP*_ = 0.9 while remaining minimally changed for the weakly adsorbed lcQDs with *χ*_*DOTAP*_ = 0.5. Figure 7D shows that the initial local densities of the sodium ions are similar for all three lcQDs because this density is defined by the DOPG content of the membrane. Similar to the chloride ions, the number of sodium ions decreases for the strongly adsorbing lcQDs with *χ*_*DOTAP*_ = 0.9 and is effectively unchanged for the lcQDs with *χ*_*DOTAP*_ = 0.5. The number of chloride and sodium ions for lcQDs with *χ*_*DOTAP*_ = 0.1 increase throughout the simulation due to the larger region that can accommodate these ions as the lcQD diffuses away from the membrane.

The displacement of counterions along with the increased adsorption strength observed for the lcQDs with *χ*_*DOTAP*_ = 0.9 is consistent with the counterion release mechanism, which is a common driving force in the complexation of oppositely charged biomolecules (*e*.*g*., protein, DNA, lipid membranes)^54^. In this mechanism, the release of counterions that were initially confined to the region between the membrane and lcQD leads to an increase in ion translational entropy, with release possible due to the opposite-charge attraction between DOTAP and DOPG lipids. This effect also can then strengthen interactions between oppositely charged lipids (*i*.*e*., DOTAP from the lipid coating and DOPG from the lipid membrane). Because this mechanism depends upon the number of counterions released and the concomitant interactions between oppositely charged lipids, it is observed only for the lcQDs with *χ*_*DOTAP*_ = 0.9 which both have high counterion densities and ample DOTAP-DOPG interactions. In contrast, the lcQDs with *χ*_*DOTAP*_ = 0.51 are instead repelled from the membrane as an alternative mechanism of reducing confinement, thereby leading to the increased local density of counterions in Figures 7C and 7D.

Our finding that counterion release explains the strong adsorption of highly charged cationic lcQDs is consistent with a previous study that showed that the binding probability of solid NPs coated with a fixed number of DOTAP molecules to DOPC:DOPS membranes increased almost 5-fold as the DOPS concentration was increased from 25% to 33%^22^. They also observed NP diffusion between bulk solvent and the membrane interface for the less anionic membrane, consistent with our observations (Figures S14-S16). Moreover, membrane binding was proposed to be a two-step process, where the NP first approaches the membrane due to electrostatic attraction between the lipid headgroups of the lipid coating and the membrane, followed by internalization initiated by lipid tail protrusions^55, 56^ and lipid exchange. This two-step mechanism is qualitatively similar to what we observed in our simulations with the exception of the lipid exchange step. However, this is attributed to the fact that we performed short simulations of only 50 ns, compared to 5 µs in this previous study, and the exchange of lipids between the lipid coating and the lipid membrane occurs at long timescales due to the fusion-like behavior during the NP internalization process. Thus, we anticipate that lipid reorganization and hydrophobic contact between lipid tails from the lipid coating and the lipid membrane could eventually initiate a fusion-like event that ultimately leads to lcQD internalization into the membrane on timescales beyond those studied in these simulations that could impact the experimentally observed lcQD fluorescence intensity in Figure 4.

## 3. Conclusions

We demonstrate a tunable methodology for functionalizing QDs with a lipid coating to control their interactions with biological membranes. By combining experimental observations with CG MD simulations, we establish mechanistic insight on the control of the lcQDs surface charge and subsequently, their cell staining efficiency by adjusting the lipid-to-particle stoichiometry during preparation. Experiments and simulations indicate that the effective surface charge of lcQDs increases with the concentration of cationic DOTAP lipids. FACS experiments with HEK293 cells show a commensurate increase in staining intensity as a function of surface charge suggesting that electrostatics, mediated by lipid ratio and compositions at preparation, play a primary driving force for lcQD–membrane interactions. Explicit solvent simulations further indicate that the release of counterions from the lcQD-membrane interface promotes strong adsorption for lcQDs with high cationic lipid contents. Overall, this work combines experiments and simulations to demonstrate a facile approach to controlling the surface charge of nanoparticles through a self-assembled lipid monolayer. This enhanced tunability of surface properties is advantageous in the design of membrane-active biocompatible inorganic nanoparticles that can act as membrane potential sensors, bioimaging agents, and drug carriers.

## 4. Materials and Methods

### Chemicals and Materials

CdSe/ZnS quantum dots (type I) coated with octadecylamine (ODA) with diameter of 5 and 11 nm, and emission maximum of 600 nm and 620 nm, respectively, were purchased form Ocean Nanotech (San Diego, USA), and a 5 nm, 540 nm emitting quantum dot (type I) coated with ODA was purchased from Creative Diagnostics (New York, USA). 1,2-dimyristoyl-sn-glycero-3-phosphocholine (DMPC), Dioleoyl-3-trimethylammonium-propane (DOTAP) were purchased from Avanti Polar Lipids (Alabaster, AL). Sodium cholate hydrate, Amberlite XAD-2, and all other materials were purchased from Sigma-Aldrich (St. Louis, MO) at the appropriate purity level.

### Preparation of lcQDs and characterization

lcQDs were formed by the hydrophobic interaction of the as-synthesized QDs’ capping ligands octadecylamine (ODA), and the lipids’ alkyl chain. Briefly, 100 µl of 1 µM QD in toluene was mixed with a DMPC:DOTAP mixture in chloroform with 180 µl mixture of 1:1:1 methanol:chloroform:DCM (Dichloromethane) with 27 µmol of sodium cholate to achieve a desired lipids:QD ratio. Organic solvents were evaporated under vacuum at 50ºC for 30 min using a benchtop vacuum concentrator (CentriVap, Labconco). The thin film formed on the glass vial walls was resuspended in 500 µl of 20 mM Tris buffer, pH 7.4. The samples were then sonicated for 1 minute in a probe sonicator at 26 W, 20kHz (VCX 130, Sonics), followed by cholate removal with Amberlite XAD-2 beads for 60 minutes at 25ºC. The beads were discarded by filtering the samples through a 0.2 µm syringe filter. Samples were characterized for hydrodynamic size by dynamic light scattering (DLS), surface charge by zeta potential measurements, and relative normalized brightness by absorption and emission spectra (see SI).

### TGA-GC/MS analysis

∼1 mg of freeze dried lcQDs was subjected to a TGA oven, with a heating rate of 20°C/min from 25 to 900°C under nitrogen atmosphere (balance purge 80 mL/min, sample purge 20 mL/min) on alumina crucibles. TGA was connected to a transfer-line (TL9000) of ‘Red Shift’ heated to 280°C, which is connected to a 100 μL loop. The gas mixture was then separated over a 5MS (30 m × 0.25 mm) column using helium gas carrier at 1.0 ml/min flow. The column temperature was raised from 40 to 300°C at 20°C/min and then held at 300°C for 5 min. The mass spectrometer conditions were as follows: electronic impact ionization at 70 eV, source temperature of 220°C, and GC transfer line at 250°C. The mass range was 35−300 Da with 0.2 s dwell time. The data were collected and analyzed by PerkinElemer GCMS TurboMass software.

### Fluorescence-Activated Cell Sorting (FACS) of lcQD-labeled cells

The degree of lcQD-membrane interaction was assessed via FACS measurements of lcQD-labeled human embryonic kidney 293 cells (HEK293).

Each lcQD construct was diluted to the same concentration, and HEK293 staining was performed by incubating 1 million cells/ml in a buffer solution (140 mM NaCl, 2.8 mM KCl, 2 mM CaCl_2_, 2 mM MgCl_2_, 10 mM glucose, and 10 mM HEPES, pH 7.4) with 3 nM of each lcQD construct for 10 minutes.at room temperature, Cells were then washed 3 times by centrifugation at 1000 RPM for 3 minutes, removing the supernatant, and resuspended in PBS buffer. Trypan Blue viability staining was performed on TC20 cell counter (BIO-RAD) prior to the lcQD incubation. FACS analysis was conducted on an LSRFortessa cell analyzer (BD Biosciences) 30 min after cell staining, using 488 nm excitation and fluorescence emission collected with a 610 nm/20 nm filter.

### Simulations of lipid self-assembly on QDs

We simulated the self-assembly of lipids on the surface of QDs using coarse-grained (CG) simulations with the Dry Martini^36^ force field. This force field builds upon the explicit solvent version of Martini^51-53, 57, 58^ by recalibrating the original interaction matrix of Lennard Jones (LJ) potentials to forego the need to model water explicitly and speed up simulations by up to two orders of magnitude. Since Dry Martini excels at reproducing lipid properties and modeling events primarily driven by the hydrophobic effect, we found it suitable to use for our lipid self-assembly protocol with large (5 nm diameter) QDs. A stochastic dynamics integrator with a timestep of 20 fs was used with the neighbor list constructed every 10 steps using the Verlet scheme and a cutoff of 1.4 nm. Van der Waals interactions were modeled using a cutoff scheme with the force switched to zero from 0.9 nm to 1.2 nm. Electrostatic interactions were modeled using a cutoff scheme with the potential shifted to zero at 1.2 nm. The dielectric constant was set to 15. Simulations were performed in the *NVT* ensemble at 300 K. The CG models for the QDs and DOTAP were taken from our previous work ^33^. DMPC was represented with the DLPC Martini model. Molecular dynamics (MD) simulations were performed with the GROMACS package, version 2023.^59^

A single QD was centered in a 50 nm × 50 nm × 50 nm box and lipids were inserted in random positions. The system was neutralized by inserting the corresponding number of counterions. Due to the stochastic nature of lipid self-assembly, convergence assessment was non-trivial; we therefore performed simulations for 3,000 ns, which was found to yield consistent behavior across multiple systems after several trial runs. Configurations were saved every 1 ns. At the end of each simulation, the lcQD was extracted for further analyses and excess lipids were removed. Four replicates were performed for each system. The total number of adsorbed lipids and the solvent-accessible surface area (SASA)^60^ were calculated and used to assess convergence (Figure S6 and S10). This procedure is schematically shown in Figure S5, and Tables S1 and S2 contain the information of the topology for each extracted lcQD and replicate. This protocol was performed for 2 nm and 5 nm diameter QDs.

### Simulations of lcQD-membrane interactions

We used the explicit solvent CG force field Martini^57^, version 2.3^51^ with the refined polarizable water model^52^ to study the interactions between positively charged lcQDs and negatively charged lipid membranes. All explicit solvent simulations were performed with a leap-frog integrator and a timestep of 20 fs. The neighbor list was constructed every 20 steps using a Verlet scheme with a cutoff of 1.225 nm. Van der Waals interactions were modeled using a cutoff scheme with the potential shifted to zero at 1.1 nm. Electrostatic interactions were modeled using the smooth Particle-Mesh-Ewald with the potential shifted from to zero at 1.1 nm, 4^th^ order interpolation, and a Fourier spacing of 0.12 nm. The dielectric constant was set to 2.5. The temperature was controlled at 300 K using the velocity-rescale thermostat coupling^61^ every 1 ps. Equilibration simulations in the *NPT* ensemble were performed with semi-isotropic pressure coupling using the Berendsen barostat coupling the pressure every 5 ps. For production simulations, we switched to the Parrinello-Rahman barostat coupling the pressure every 12 ps. MD simulations were performed with the GROMACS package, version 2023.^59^

Out of the four replicates performed for the self-assembly simulations with *χ*_*DOTAP*_ = 0.5, the replicate that showed full encapsulation with the largest number of adsorbed lipids was selected for these sets of simulations. The replicates selected from the simulations with *χ*_*DOTAP*_ of 0.1 and 0.9 were the ones that showed full encapsulation and the number of adsorbed lipids closest to that of the 50% DOTAP lcQD. We solvated the extracted 2 nm diameter lcQDs with an 18 nm × 18 nm membrane (774 DOPC and 192 DOPG) in a 24 nm tall box with approximately 50,000 polarizable water beads^52^, 150 mM NaCl, and counterions to neutralize the charged lipid coating. For each replicate, the lipid membrane was initialized randomly. The lcQD was randomly oriented and positioned so that the minimum distance between the surface of the lipid coating and the surface of the membrane was approximately 2 nm. The three coordinates of all beads that compose the QD core were restrained with a 1,000 kJ·mol^-1^·nm^-2^ during equilibration simulations to prevent translation. The energy of the system was minimized and followed by a 1 ns *NVT* simulation to equilibrate the solvent. Subsequently, a 10 ns *NPT* simulation was performed to allow the lipid membrane to relax. After equilibration, the position restraints were released and a 50 ns production simulation was performed, sampling every 50 ps. A total of 16 independent replicates were performed for each of the lcQDs with*χ*_*DOTAP*_ of 0.1, 0.5, and 0.9 with each replicate having unique initial configurations and velocities.

## Data availability

Raw and processed data for the main simulation and experimental data presented in this work is available at Dryad: 10.5061/dryad.ns1rn8q7c.

## Acknowledgements

This work has received funding from the European Research Council (ERC) under the European Union’s Horizon 2020 research and innovation program under grant agreement No. 669941 and ERC-POC grant agreement No. 779896, by the BER program of the Department of Energy Office of Science grant DE-SC0020338, by the STROBE National Science Foundation Science & Technology Center, Grant No. DMR-1548924, by the Israel Science Foundation grant # 813/19, and by the Bar-Ilan Research & Development Co, the Israel Innovation Authority, Grant No. 63392. RCV was supported by the National Science Foundation under the CAREER Award No. DMR-2044997. CAHZ and RCV acknowledge the support provided by the Graduate Engineering Research Scholars–Advanced Opportunity Fellowship from the University of Wisconsin– Madison, the Chemistry-Biology Interface Training Program (T32GM152341), funded by the National Institute of General Medicine Sciences, and the National Science Foundation through a Graduate Research Fellowship. This work used TAMU FASTER resources at Texas A&M University through allocation CHM230045 from the Advanced Cyberinfrastructure Coordination Ecosystem: Services & Support (ACCESS)^62^, which is supported by the National Science Foundation under the Office of Advanced Cyberinfrastructure awards #2138259, #2138286, #2138307, #2137603, and #2138296. We would like to thank Hagit Hauschner for contributing with FACS experiments in Figure 4.

## Notes

https://doi.org/10.5061/dryad.ns1rn8q7c

### Competing Interest Statement

The authors have declared no competing interest.

