## Supplementary Information for "Tunable electrostatic interactions of lipid-coated quantum dots with biological membranes"

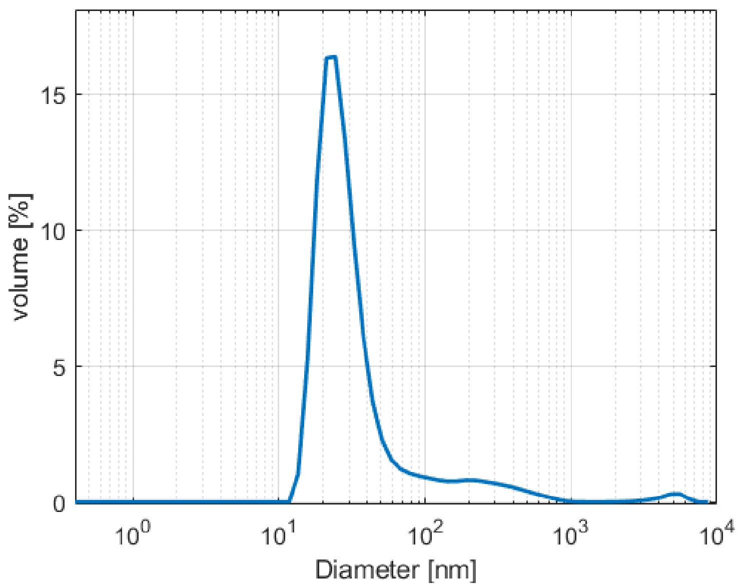


Figure S1. DLS size distribution by volume of typical lcQDs preparation. The hydrodynamic diameter of lcQD construct of DMPC:DOTAP 1:1, 3500 lipids per particle, 33.98 ± 9.83 nm by intensity, 92% of sample volume.

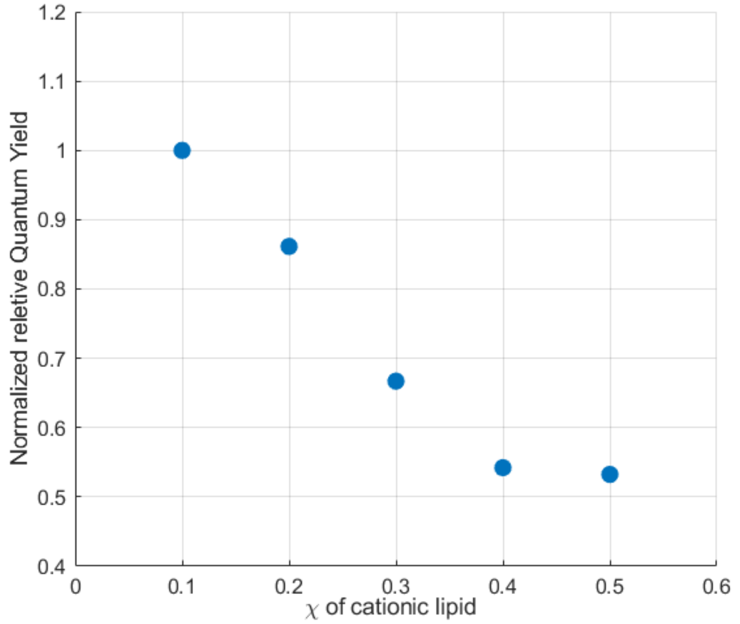


Figure S2. Normalized relative brightness of lcQDs constructs of experiments in figure 1. Total emission intensity divided by the O.D. and normalized relatively to each other's. The QDs emission is decreased with the increase adsorption of charged lipids.

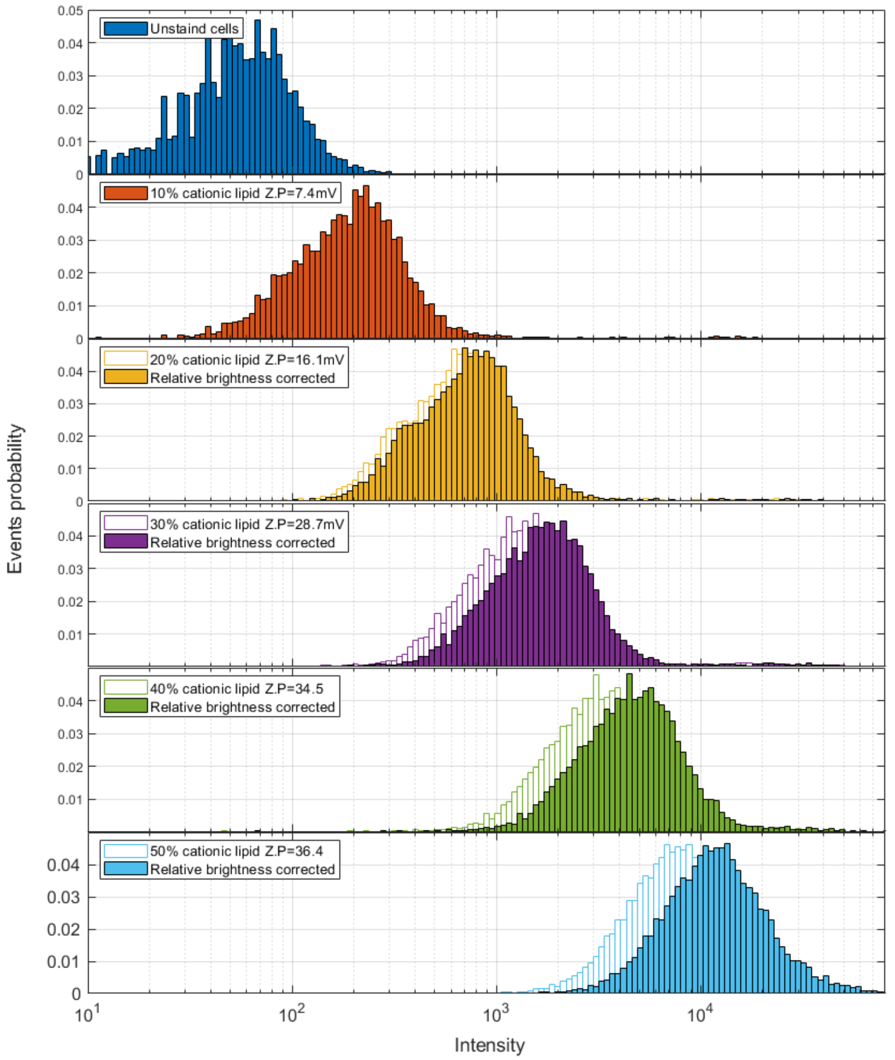


Figure S3. Flow cytometry results for HEK cells stained with the same series of 11 nm lcQDs from S2 with varying DOTAP/total lipids ratios adjusted to relative brightness.

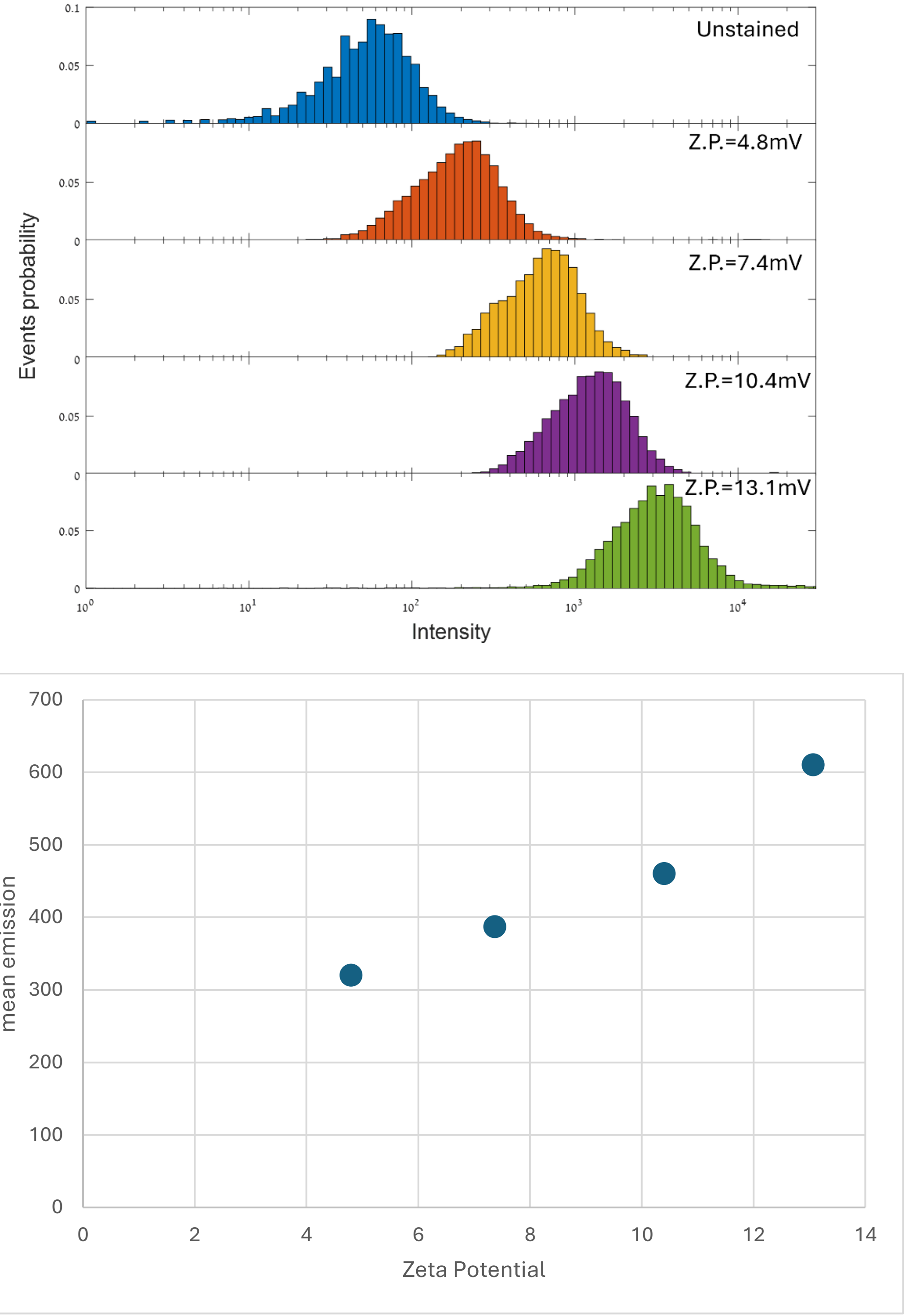


Figure S4**:** (A) Flow cytometry results for HEK cells stained with a series of 5 nm lcQDs with varying DOTAP/total lipids. Cell staining is surface charge dependent, staining intensity increases with particle's charge. (B) Corresponding intensity measurement for QDs used in (A) as function of lcQDs' zeta potential.

### Supplementary Methods

#### Dynamic light scattering:

Dynamic light scattering data were collected using the Zetasizer Nano ZS instrument (Malvern Instruments, Great Britain) operating at 25°C with a laser wavelength λ of 620 nm and a scattering angel of θ 173°. Measurements were made using polystyrene low-volume microcuvettes (BrandTech Scientific, Inc. USA). Cuvettes were filled with 100 μL of lcQDs dispersion for DLS characterization. Exemplary DLS data is shown in Figure S1.

#### Zeta Potential

The zeta potential measurements were performed using the electrophoretic light scattering technique on a Zetasizer Nano ZS analyzer (Malvern Instruments, Great Britain). using laser to determine electrophoretic mobility. The measurements were performed in a high concentration cell with palladium electrodes at 25°C and pH 7.4 in a Tris buffered solution containing no chlorine ions. The results were processed using the Dispersion Technology Software 6.2 (Malvern Instruments).

#### Absorbance spectroscopy:

The absorbance spectrums of NPs were obtained with Ocean Optics USB4000 spectrophotometer. The spectra of NPs were obtained at a scan rate of 1200 nm/min with 1 cm path length in a quartz cuvette.

#### Fluorescence measurement

The emission spectra of NPs were obtained by Ocean Optics USB4000 spectrophotometer (Ostfildern, Germany) at a scan rate of 1200 nm/min with 1 cm path length, in a quartz cuvette, with excitation and emission slits widths 5 nm. Samples were excited using a 380 nm wavelength light.

#### HEK293 cells line maintenance

Human Embryonic Kidney 293 (HEK293) cells were maintained under standard growth conditions in a humidified incubator in high glucose Dulbecco's modified Eagle's medium supplemented with 10% fetal calf serum, 2 mM glutamine, 100 units/mL penicillin G, and 100 μg/mL streptomycin (Beit HaEmek, Biological Industries, Israel) at 37°C and 5% $CO_2$.

**<u>Simulations of lipid-self assembly</u>**

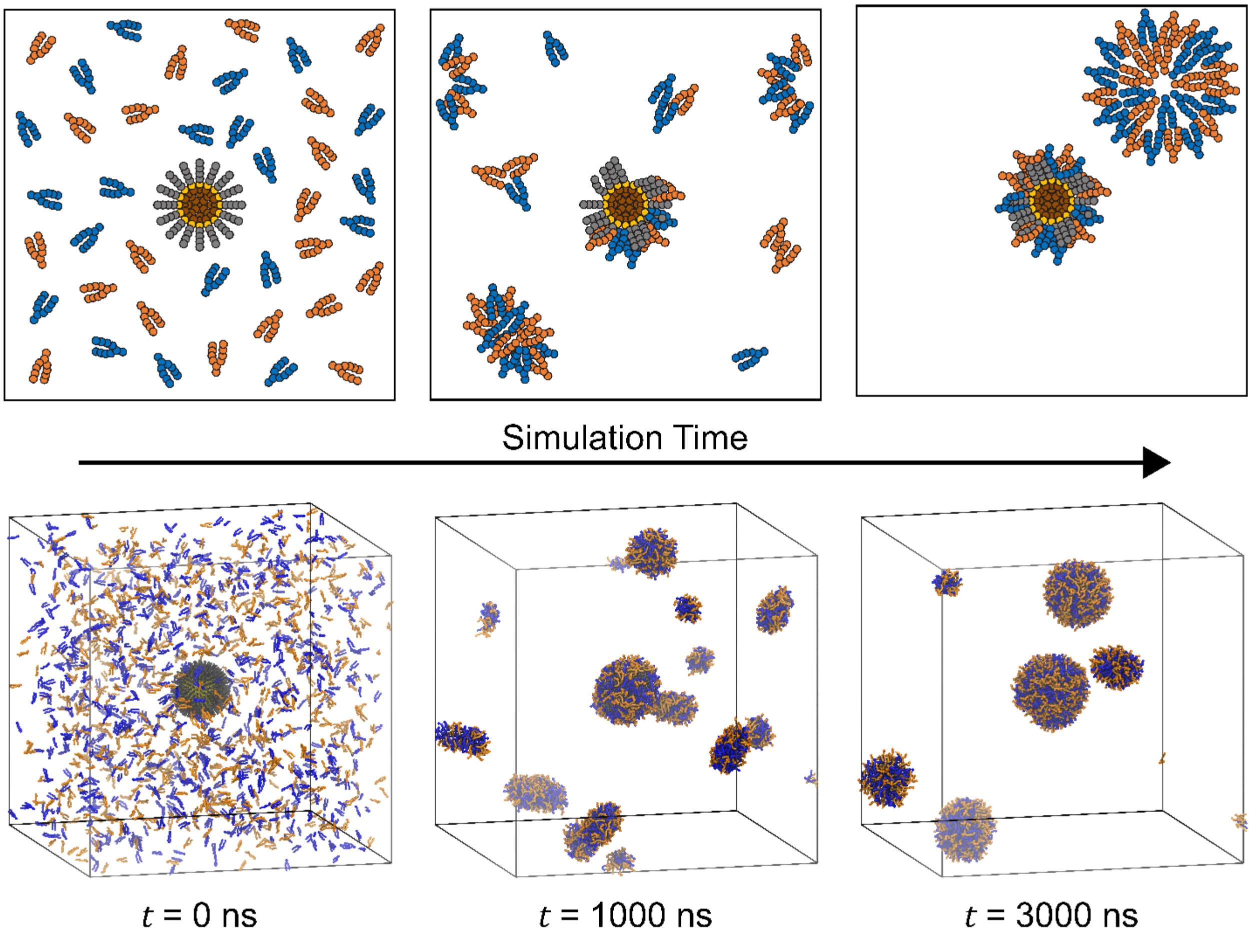


Figure S5. Schematic of lipid self-assembly on single quantum dot. Counterions are present but omitted from the schematic and simulation snapshots for visual purposes. Blue corresponds to cationic lipids (DOTAP) and orange corresponds to zwitterionic lipids (DMPC). Simulation snapshots correspond to a single replicate of the 5 nm diameter quantum dot.

Table S1. Details of extracted lipid-coated quantum dots (QDs) after lipid self-assembly from simulations in which encapsulation efficiency was assessed.

| QD diameter | Number of total lipids | Replicate number | Number of DLPC | Number of DOTAP |
|---|---|---|---|---|
| 5 nm | 500 | 1 | 68 | 65 |
| | | 2 | 92 | 97 |
| | | 3 | 39 | 48 |
| | | 4 | 69 | 68 |
| | 1000 | 1 | 98 | 109 |
| | | 2 | 199 | 200 |
| | | 3 | 199 | 187 |
| | | 4 | 34 | 32 |
| | 1500 | 1 | 156 | 148 |
| | | 2 | 215 | 243 |
| | | 3 | 41 | 39 |
| | | 4 | 32 | 42 |
| | 2000 | 1 | 136 | 148 |
| | | 2 | 209 | 213 |
| | | 3 | 305 | 325 |
| | | 4 | 337 | 340 |
| | 2500 | 1 | 44 | 60 |
| | | 2 | 429 | 373 |
| | | 3 | 334 | 365 |
| | | 4 | 348 | 356 |
| | 3000 | 1 | 117 | 117 |
| | | 2 | 227 | 223 |
| | | 3 | 350 | 381 |
| | | 4 | 213 | 191 |
| | 3500 | 1 | 242 | 229 |
| | | 2 | 425 | 407 |
| | | 3 | 462 | 427 |
| | | 4 | 952 | 940 |
| | 4000 | 1 | 193 | 167 |
| | | 2 | 329 | 320 |
| | | 3 | 232 | 229 |
| | | 4 | 233 | 253 |
| 2 nm | 500 | 1 | 53 | 41 |
| | | 2 | 56 | 45 |
| | | 3 | 17 | 21 |
| | | 4 | 171 | 169 |
| | 1000 | 1 | 37 | 47 |
| | | 2 | 217 | 201 |
| | | 3 | 146 | 165 |
| | | 4 | 263 | 255 |
| | 1500 | 1 | 101 | 90 |
| | | 2 | 192 | 178 |
| | | 3 | 333 | 337 |
| | | 4 | 402 | 421 |

Table S2. Details of extracted lipid-coated quantum dots (QDs) after lipid self-assembly from simulations in which the influence of initial DOTAP concentration ($\chi_{DOTAP}^{overall}$) was assessed.

| QD diameter | $\chi_{DOTAP,overall}$ | Replicate number | Number of DLPC | Number of DOTAP |
|---|---|---|---|---|
| 5 nm | 0.10 | 1 | 135 | 20 |
| | | 2 | 396 | 52 |
| | | 3 | 190 | 16 |
| | | 4 | 84 | 11 |
| | 0.20 | 1 | 162 | 34 |
| | | 2 | 52 | 10 |
| | | 3 | 234 | 61 |
| | | 4 | 296 | 82 |
| | 0.30 | 1 | 96 | 43 |
| | | 2 | 265 | 122 |
| | | 3 | 450 | 180 |
| | | 4 | 212 | 67 |
| | 0.40 | 1 | 285 | 183 |
| | | 2 | 168 | 102 |
| | | 3 | 319 | 225 |
| | | 4 | 105 | 70 |
| | 0.50 | 1 | 156 | 148 |
| | | 2 | 215 | 243 |
| | | 3 | 41 | 39 |
| | | 4 | 32 | 42 |
| | 0.60 | 1 | 87 | 139 |
| | | 2 | 104 | 181 |
| | | 3 | 196 | 320 |
| | | 4 | 59 | 90 |
| | 0.70 | 1 | 108 | 224 |
| | | 2 | 54 | 136 |
| | | 3 | 29 | 69 |
| | | 4 | 53 | 107 |
| | 0.80 | 1 | 36 | 193 |
| | | 2 | 17 | 84 |
| | | 3 | 97 | 408 |
| | | 4 | 51 | 153 |
| | 0.90 | 1 | 44 | 295 |
| | | 2 | 46 | 324 |
| | | 3 | 35 | 348 |
| | | 4 | 53 | 546 |
| 2 nm | 0.10 | 1 | 105 | 10 |
| | | 2 | 151 | 21 |
| | | 3 | 107 | 8 |
| | | 4 | 188 | 21 |
| | 0.50 | 1 | 53 | 41 |
| | | 2 | 56 | 45 |
| | | 3 | 17 | 21 |
| | | 4 | 171 | 169 |
| | 0.90 | 1 | 19 | 147 |
| | | 2 | 13 | 135 |
| | | 3 | 29 | 291 |
| | | 4 | 16 | 126 |

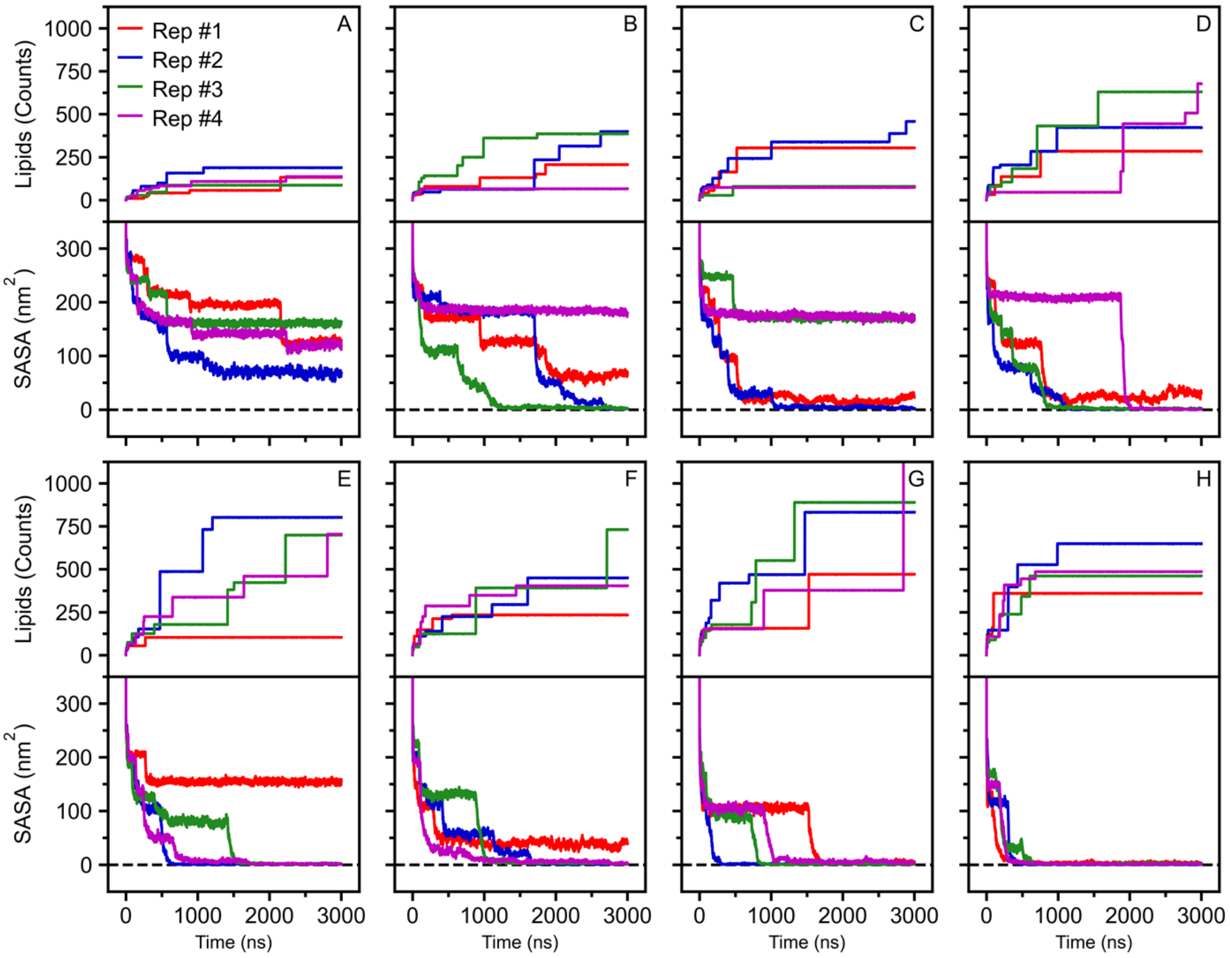


Figure S6. Convergence of lipid self-assembly on 5 nm quantum dot. Lipid count and solvent accessible surface area (SASA) were plotted for each of the four replicates for total lipids: (A) 500, (B) 1000, (C) 1500, (D) 2000, (E) 2500, (F) 3000, (G) 3500, (H) 4000. The *gmx sasa*[1] tool was used to compute the SASA with a probe radius of 0.26 nm and 4800 grid points.

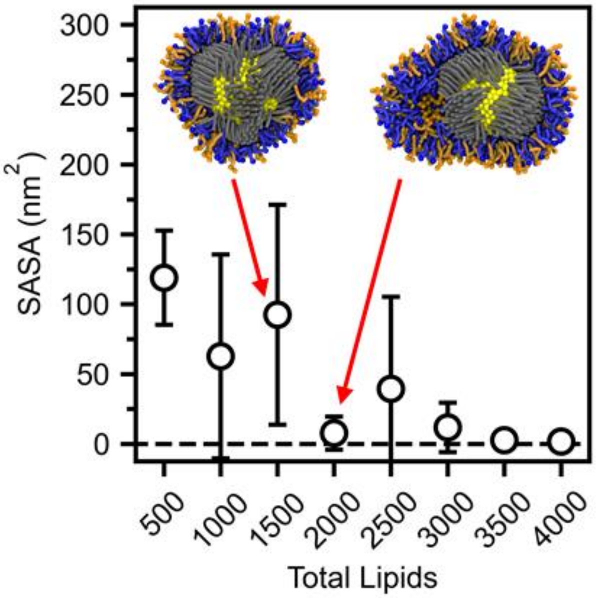


Figure S7. Assessment of encapsulation efficiency. 1500 total lipids was chosen as full encapsulation since it resulted in fewer multilayer lipid structures compared to the 2000 total lipids simulations. Error was computed as the standard deviation across four replicates. Blue corresponds to cationic lipids (DOTAP) and orange corresponds to zwitterionic lipids (DMPC).

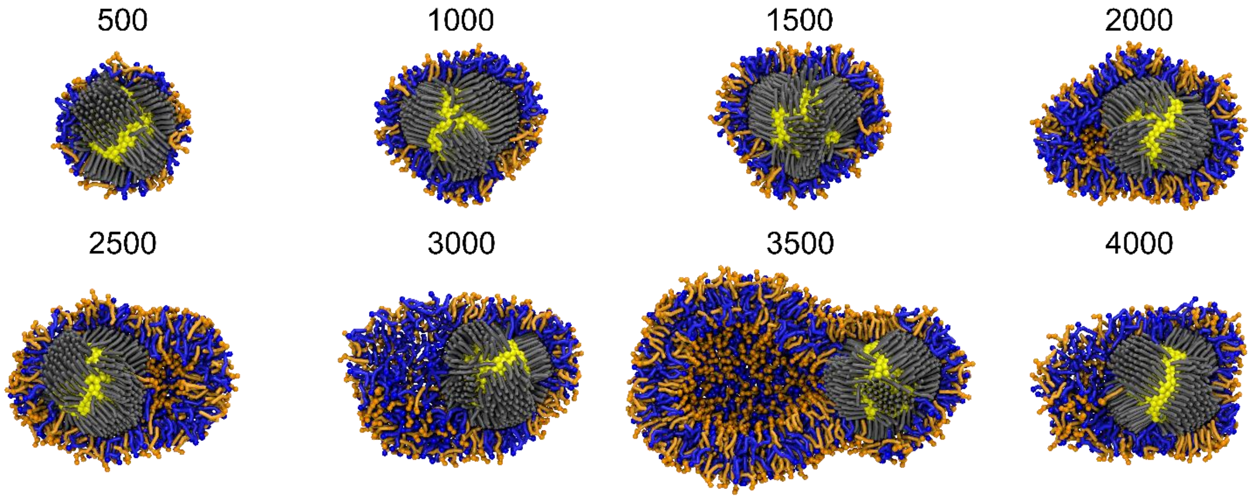


Figure S8. Cross-sectional view of final lipid-coated quantum dots from simulations in which encapsulation efficiency was assessed. Blue corresponds to cationic lipids (DOTAP) and orange corresponds to zwitterionic lipids (DMPC).

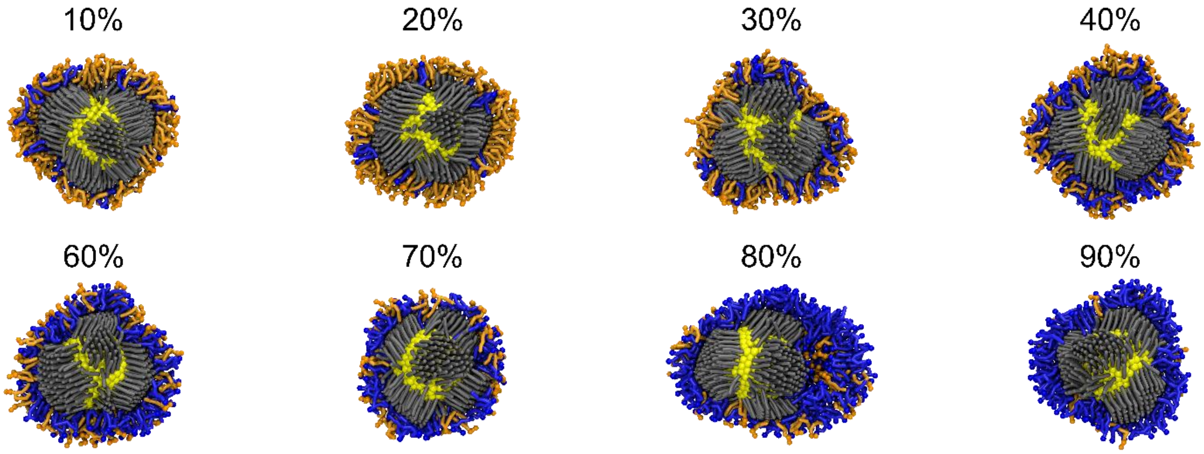


Figure S9. Cross-sectional view of final lipid-coated quantum dots from simulations in which the influence of initial DOTAP concentration (%) was assessed. Blue corresponds to cationic lipids (DOTAP) and orange corresponds to zwitterionic lipids (DMPC).

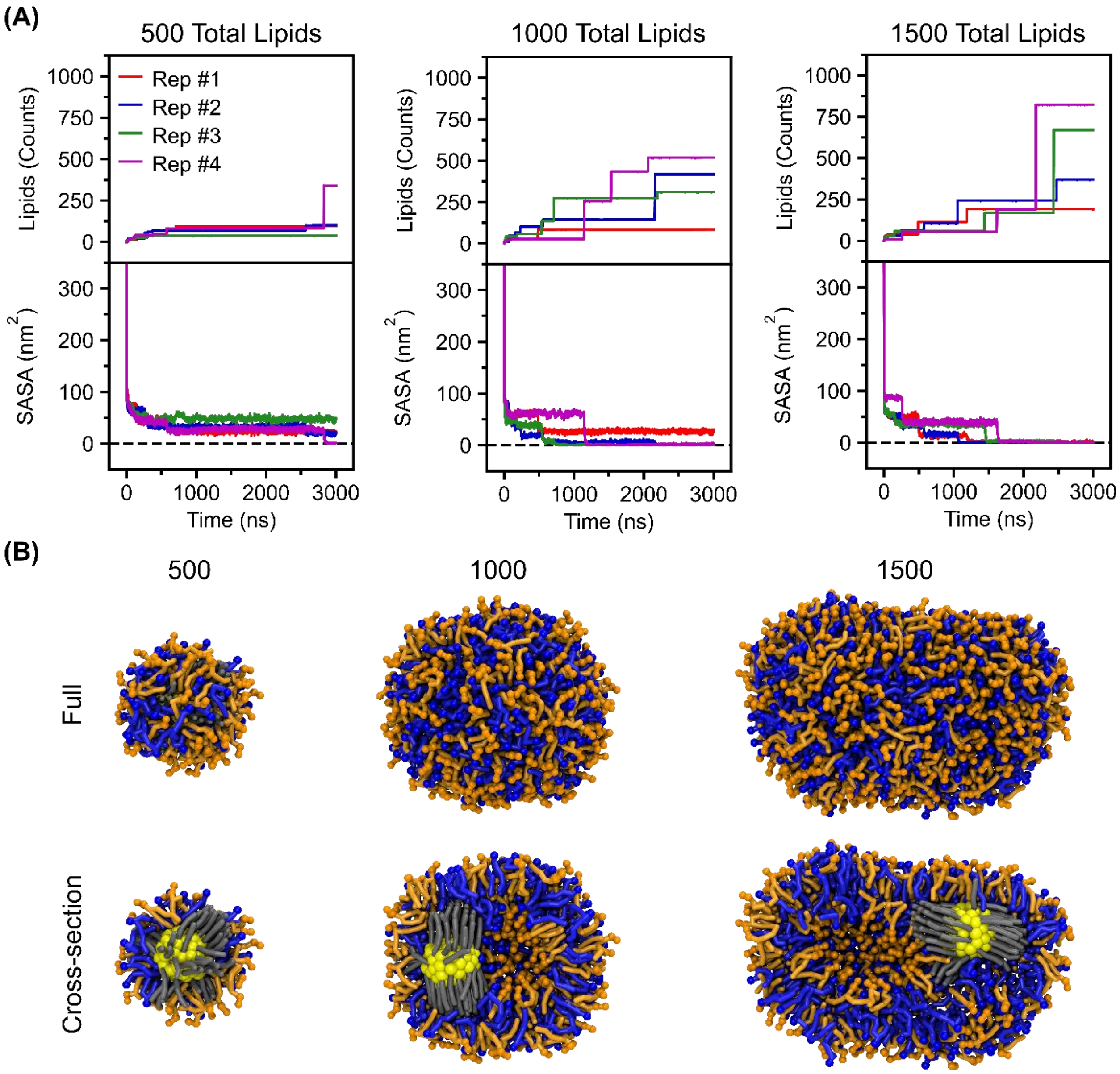


Figure S10. Convergence of lipid self-assembly on 2 nm quantum dot. (A) Lipid count and solvent accessible surface area (SASA) were plotted for each of the four replicates. The *gmx sasa*[1] tool was used to compute the SASA with a probe radius of 0.26 nm and 4800 grid points. (B) Full and cross-sectional views of final lipid-coated quantum dots from simulations in which encapsulation efficiency was assessed. Blue corresponds to cationic lipids (DOTAP) and orange corresponds to zwitterionic lipids (DMPC).

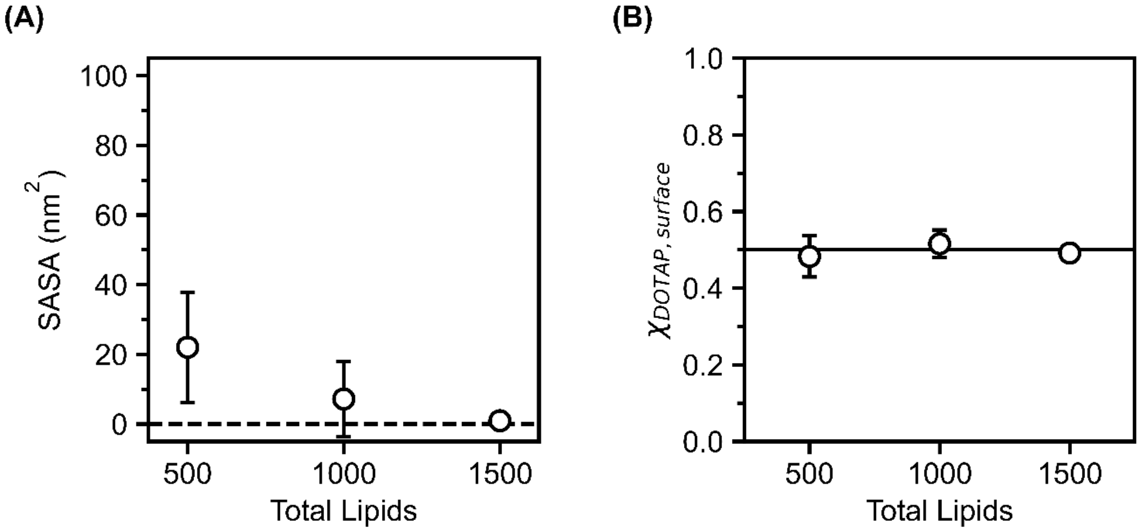


Figure S11. (A) Assessment of encapsulation efficiency. (B) Concentration of DOTAP on the surface lipid coating ($\chi_{DOTAP,surface}$) as a function of total lipids. The error in (A) and (B) was computed as the standard deviation across four replicates.

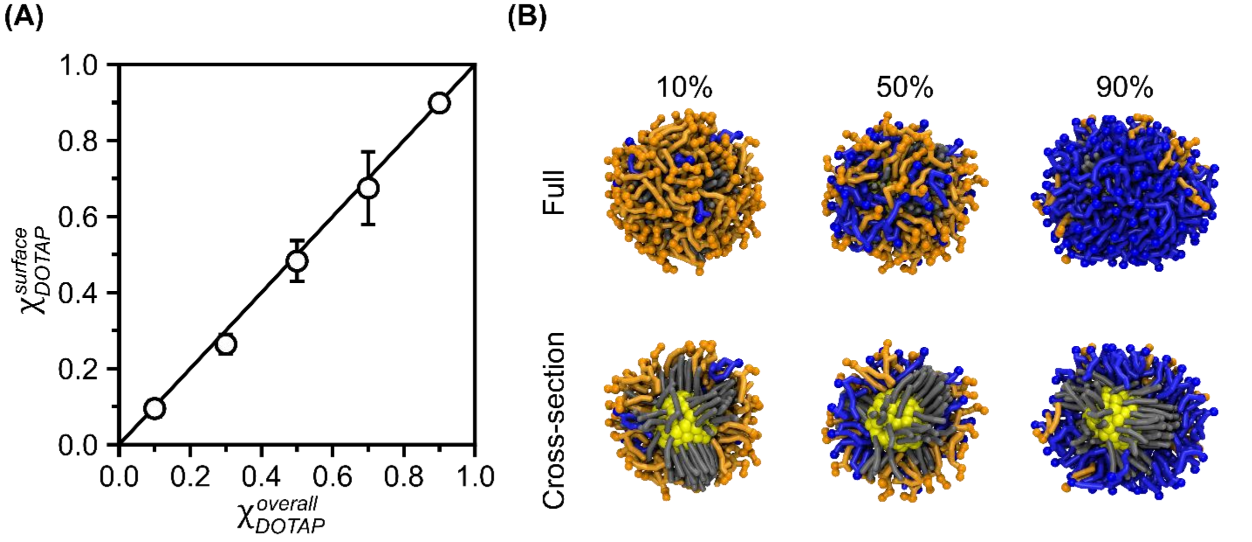


Figure S12. (A) Concentration of DOTAP on the surface lipid coating ($\chi_{DOTAP}^{surface}$) as a function of the total DOTAP concentration in the simulation box ($\chi_{DOTAP}^{overall}$). Error was computed as the standard deviation across four replicates. (B) Full and cross-sectional views of final lipid-coated quantum dots from simulations in which the influence of initial DOTAP concentration (%) was assessed. Blue corresponds to cationic lipids (DOTAP) and orange corresponds to zwitterionic lipids (DMPC).

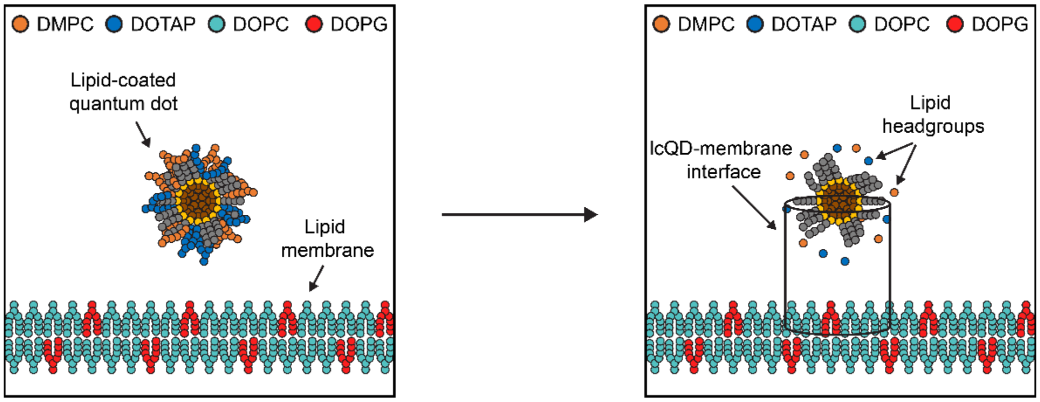


Figure S13. Definition of lcQD-membrane interface. An effective radius centered on the quantum dot is defined using the headgroups of the lipids within the lipid coating. The interface is defined as the cylindrical region with this effective radius from the center of mass of the quantum dot and the midplane of the lipid membrane.

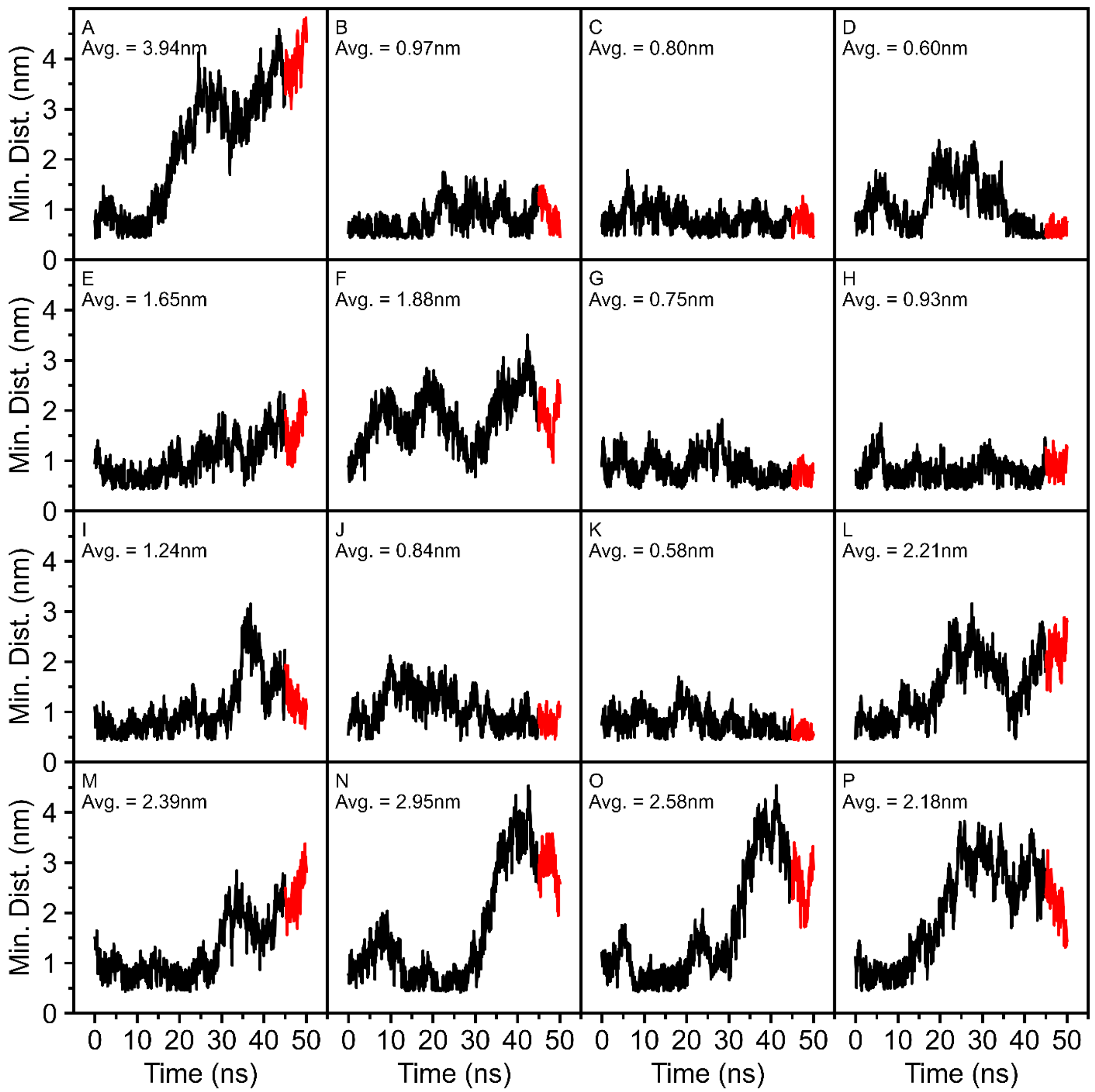


Figure S14. Minimum distance between the lipid-coated quantum dot with $\chi_{DOTAP,surface}$ of 10% and the lipid membrane. Calculation was performed for all 16 replicates using the *gmx mindist* tool between any bead from the lipid-coated quantum dot (monolayer lipids and ligands) and any bead from the lipid membrane. The data colored in red correspond to the last 5 ns of the simulation which were used to assess adsorption strength; the average value of these data is included in each subplot.

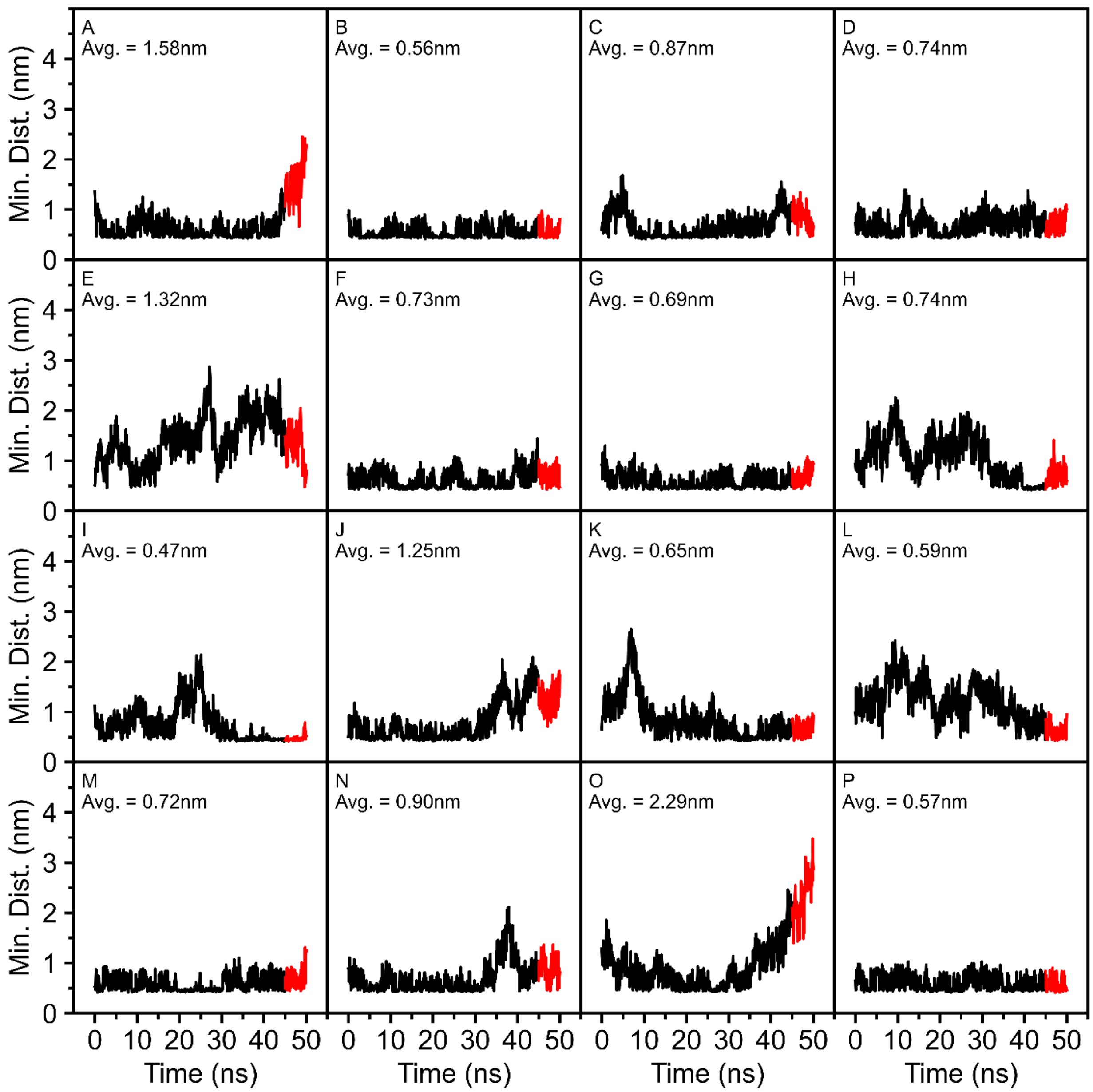


Figure S15. Minimum distance between the lipid-coated quantum dot with $\chi_{DOTAP,surface}$ of 50% and the lipid membrane. Calculation was performed for all 16 replicates using the *gmx mindist* tool between any bead from the lipid-coated quantum dot (monolayer lipids and ligands) and any bead from the lipid membrane. The data colored in red correspond to the last 5 ns of the simulation which were used to assess adsorption strength; the average value of these data is included in each subplot.

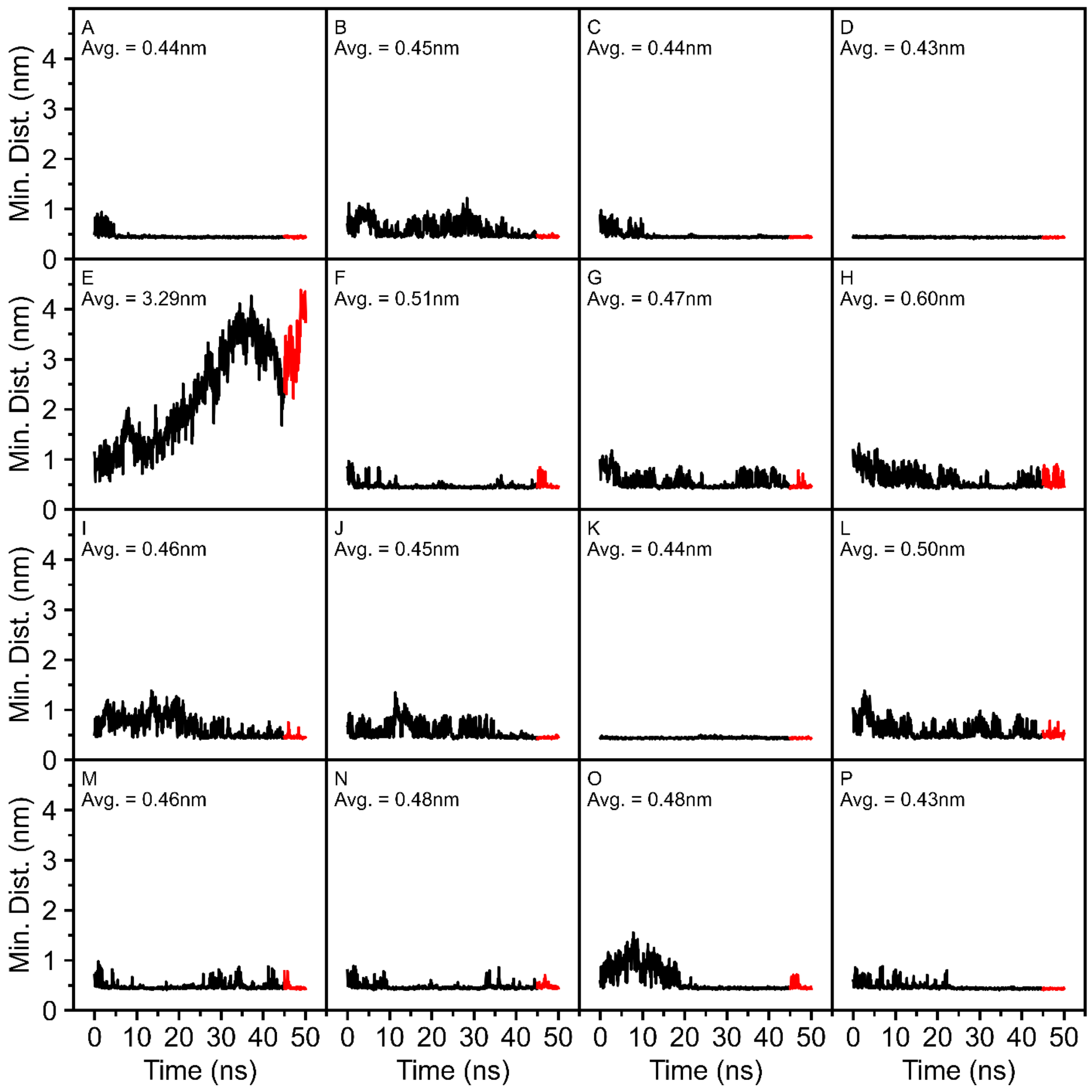


Figure S16. Minimum distance between the lipid-coated quantum dot with $\chi_{DOTAP,surface}$ of 90% and the lipid membrane. Calculation was performed for all 16 replicates using the *gmx mindist* tool between any bead from the lipid-coated quantum dot (monolayer lipids and ligands) and any bead from the lipid membrane. The data colored in red correspond to the last 5 ns of the simulation which were used to assess adsorption strength; the average value of these data is included in each subplot.
